# *In silico* identification of novel peptides as potential modulators of Aβ42 Amyloidogenesis

**DOI:** 10.1101/2022.04.20.488983

**Authors:** Kavita Kundal, Santhosh Paramasivam, Amit Mitra, Nandini Sarkar

## Abstract

Alzheimer’s Disease is a neurodegenerative disease for which no cure is available at present. The presence of amyloid plaques in the extracellular space of neural cells is the key feature of this fatal disease. Amyloid-Beta (Aβ) is a 40-42 amino acid peptide and the main component of amyloid plaques. This peptide is produced by the proteolysis of Amyloid Precursor Protein by presenilin. Deposition of 42 residual Aβ peptides forms fibrils structure, leading to disruption of neuron synaptic transmission, inducing neural cell toxicity, ultimately leading to neuron death. To modulate the amyloidosis of Aβ peptides, various novel peptides have been investigated via molecular docking and molecular dynamic simulation studies. The sequence-based peptides were designed and investigated for their interaction with Aβ42 monomer and fibril using the molecular docking method, and their influence on the structural stability of target proteins was studied using molecular simulations. According to the docking results, amongst all the synthetic peptides, the peptide YRIGY (P6) has the highest binding affinity with Aβ42 fibril, and the peptide DKAPFF (P12) shows better binding with Aβ42 monomer. Moreover, simulation results also suggest that the higher the binding affinity, the better the inhibitory action. From these findings, it is suggested that both the peptides can modulate the amyloidogenesis, but peptide (P6) has better potential for the disaggregation of the fibrils, whereas peptide P12 stabilizes the native structure of the Aβ42 monomer more effectively and hence can serve as a potential amyloid inhibitor. Thus, these peptides can be explored as therapeutic agents against Alzheimer’s Disease.

## INTRODUCTION

Proteins are informational macromolecules, a polymer of amino acids. It should remain in a proper 3D conformation for proper functioning. Misfolding/ unfolding of proteins leads to the formation of insoluble protein aggregates termed amyloid [1]. These protein aggregates convert soluble protein into insoluble oligomers, followed by the formation of amyloid fibrils, which are associated with numerous amyloidosis diseases such as Alzheimer’s Diseases, Parkinson’s Diseases, Huntington’s Diseases, Type-2 diabetes, etc[2]. Alzheimer’s Disease is characterized by the formation of insoluble inclusions of amyloid-β protein as senile plaques in extracellular space between neurons as well as by the deposition of tau proteins as neurofibrillary tangles in the cell body of neurons [3]. The proteolytic product of APP, i.e., Aβ42 is responsible for the formation of amyloid plaques [4]. Proteolytic digestion of APP by secretase form different products as mentioned in figure 1 from which resulting Aβ peptides are responsible for AD. The mutation in genes such as APP, PS1, and PS2 [5] enhances the production of Aβ42 peptides, which further increases the Aβ42/Aβ40 ratio. This enhanced production of Aβ42 relatively increases the oligomerization and fibrillation rate [6]. Missense mutation on APP, specifically on the Aβ region, increases the tendency of peptide aggregation and the stability of oligomers [7]. Lei *et al*. stated that the aggregates generated by the coexistence of Aβ40 and Aβ42 in extracellular space consist of three varieties: Aβ42 alone, Aβ40 alone, and Aβ42+ Aβ40 mixture, and deposition of Aβ40 and the mixture of both peptides are very slow to form plaques than Aβ42 alone [8]. A higher concentration of Aβ42 in a local environment and some cell components like lipids [9] promote the aggregation of Aβ42 in amyloid plaque. As a product of a metabolic process, Aβ in monomeric form adopts random coil conformation in a normal aqueous solution. Under certain changes in physiological conditions (such as mutation, decrease in proteases efficiency, extremes of pH and temperature [10], macromolecular crowding, metal ions [11][12], etc.), monomeric form starts converting into insoluble aggregates resulting in the formation of amyloids. The mechanism behind the conformational conversion is not yet properly discovered[13]. Self-assembly of Aβ into amyloid fibrils is an exclusively dynamic process that results in the formation of different fibril structures which differ in size, stoichiometry, complexity, and shape. Researches on Aβ toxicity suggested that protofibrils/oligomers are much more toxic than the mature fibrils and lead to synaptic plasticity and neurodegeneration, whereas mature fibrils contribute to neural damage by the activation of microglia upon interaction [14][15]. In the whole sequence, some key regions/residues are critical for the nucleation of β-rich structure. In the case of Aβ42, (1DAEFRHDSGYEVHHQKLVFFAEDVGSNKGAIIGLM VGGVVIA42) three regions, 13-16 (HHQK)[16], central hydrophobic region 17-21(LVFFA), and terminal hydrophobic region 30-42(IIGLMVGGVVIA) initiate the nucleation process for the fibril formation [17][18].

**Figure 1:**
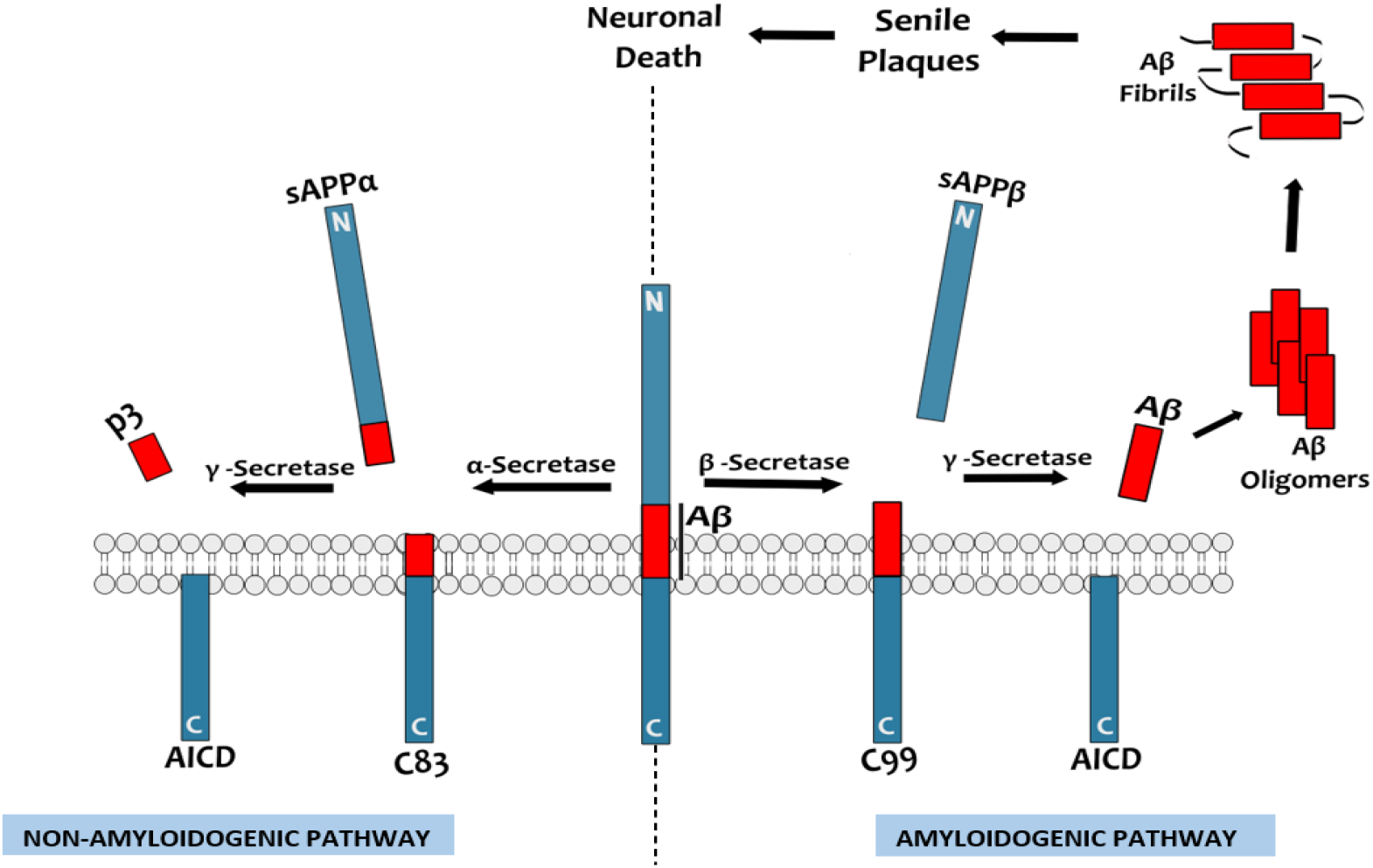
Schematic representation of APP Proteolytic Processing-APP proteolysis proceeds in two pathways: When APP is cleaved by β-secretase followed by γ -secretase, it proceeds along the amyloidogenic pathway; however, when APP is cleaved by α-secretase instead of β-secretase, it proceeds along the non-amyloidogenic pathway.

Various physical, chemical, and biomolecule-based techniques have been discovered to inhibit amyloid formation, but due to the great structural diversity of Aβ oligomers, it is a key challenge to discover a highly efficient inhibitor to prevent the formation of amyloid, which also destabilized the preformed fibrils. These strategies are categorized based on how they prevent/inhibit fibril structure formation, such as 1. Binding of small molecules to the native structure of a protein, 2. Binding of small molecules to an aggregation-prone region of a protein, 3. Binding of small molecules to the self-assembled protein and disrupting their aggregated structure 4. Peptide-based inhibitors, 5. Antibody-mediated inhibition [19].

Among all the inhibitors, peptides aroused wide interest in terms of efficiency, selectivity, better penetration, low production cost, and reduced off-targets toxicities with an additional advantage of mimicking native interaction interface. Peptides can be degraded into simple components like amino acids without leading to toxic metabolites in which D-type amino acids are more restricted for degradation [20]. These peptides were designed in such a way that it binds to amyloidogenic regions on Aβ peptides, and thereby inhibit amyloid aggregation. These inhibitory peptides can also efficiently bind to and destabilize the preformed amyloid fibril structure. Some reported peptides KLVFF[21], LPFFD [22], LVFFA[23], LPYFDa[24], iAβ5p [25], etc., were proved to have an inhibitory effect against amyloid-beta oligomers, fibrils, and its induced toxicity.

In this study, we designed a small library of peptide inhibitors against Aβ monomer (PDB ID: 1IYT) and fibrillar structure (PDB ID: 2MXU). These peptides are designed based on homologous amyloidogenic region (16K-21A) by one/two-point mutation [19] and based on structural similarity with curcumin as reported by Orjuela *et al*. [26]. These peptides are modified by terminal modifications (N-terminal Acetylation and C-terminal N-methylation), and by doing so, their inhibition efficiency increases gains stability against enzymatic degradation and improves hydrophobicity [27]. Computational tools, such as molecular docking (using the Schrodinger suite), are used for interpreting the binding affinity and efficacy of peptides against Aβ -monomers and fibril structures, and sufficient knowledge of atomic and molecular interactions can be gained. The peptide showing the highest negative binding energy, compared to controls, is considered as the best inhibitor, which was further investigated for the influence on structural stability by applying multiple nanoseconds Molecular Dynamics Simulation with Charmm36 Force Field [28] and SPC water model [29].

## MATERIALS AND METHODS

The crystal structure of monomeric and fibrillar Aβ42 was obtained from RCSB Protein Data Bank, having PDB ID of 1ITY & 2MXU, respectively. Aβ42 monomer consisting of two helical stretches of 7-26 & 28-40 residues as shown in figure-2a and Aβ42 fibril structure consists of 12 chains (A→L) with coiled and β-strand structure (fig 2b). In the fibrillar structure, maximum residues are buried inside the hydrophobic core, which helps in inducing aggregation and results in senile plaques, as shown in figure-2b. Some bioinformatics tools and databases were used to study the interactive and inhibitory properties of ligands against protein computationally, including Protein Data Bank for protein structure retrieval, the Schrodinger suite for peptide docking [30], PyMOL [31], and VMD for visualization [32], and Gromacs for MD simulations [33].

**Figure 2:**
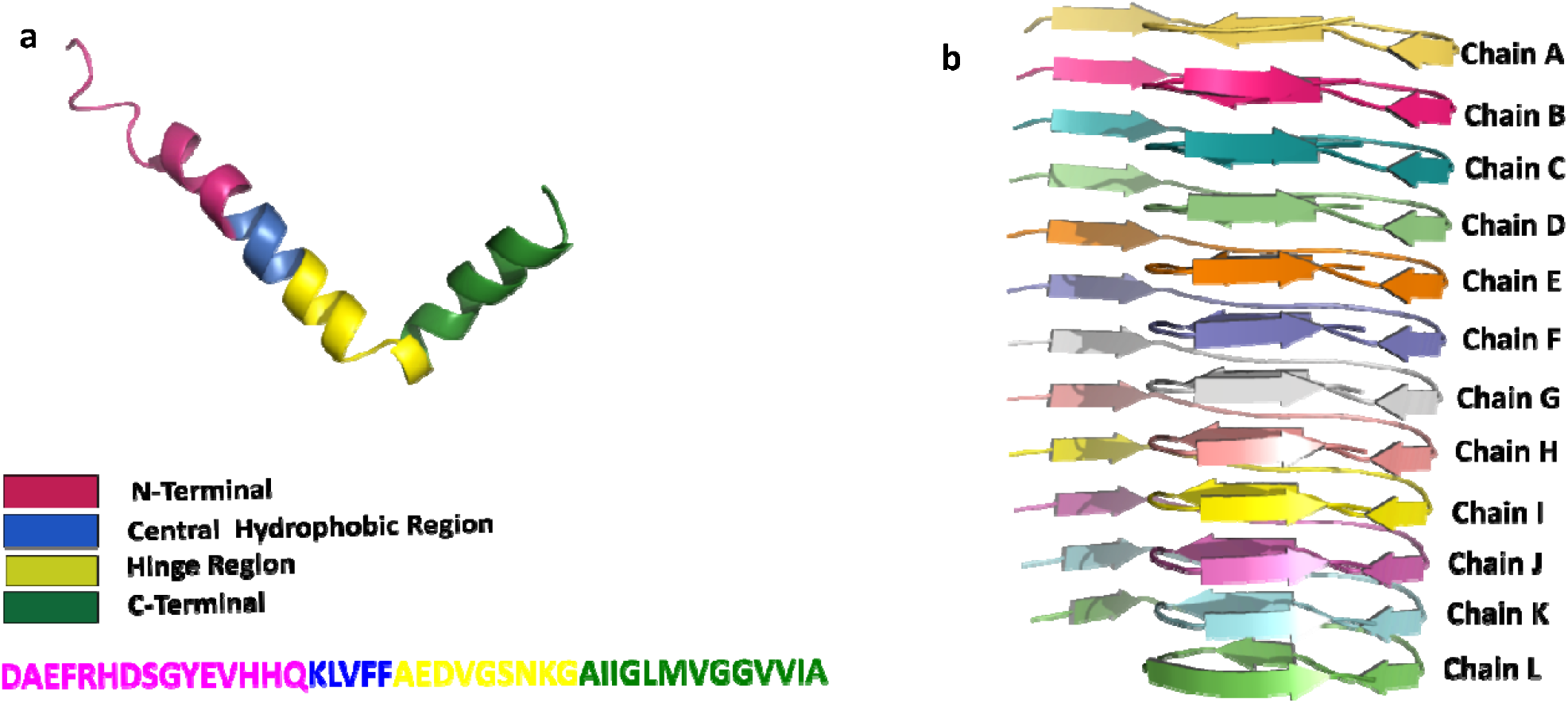
Ribbon structure of (a) Amyloid-beta (Aβ42) Peptide (PDB ID-1IYT) and (b) 12mer Amyloid-beta fibrils (PDB ID-2MXU)

### 1. Molecular Docking

Docking is a molecular technique that determines the interaction between protein and small molecules (ligands, nanoparticles, peptides, etc.)[34]. The docking study was carried out using the MM-GBSA method of MAESTRO BioLuminate with an OPLS2005 force field and VSGB2.0 solvent model for measuring the difference in the binding energy of protein (1IYT & 2MXU) with different peptide inhibitors. The Biologics suite offers a variety of options for protein modeling, protein analysis, peptide docking, and protein engineering [35][36][37]. Proteins (1IYT & 2MXU) were downloaded from PDB and prepared using Protein Preparation Wizard in the Schrodinger Suite 2020.2 release, MAESTRO BioLuminate. The protein preparation involved the assignment of bond orders, the addition of hydrogens, deletion of water beyond 5Aº, optimization, and minimization. Identification of the best top-ranked receptor binding sites of proteins was carried out by the SiteMap tool. The highly potential binding pocket was chosen based on D-score (should be in 0.9 ≤.x≥1.5) as listed in table1. Peptide docking is used for MM-GBSA execution as it is the most accurate method for binding energy estimation. Within the Peptide docking application, a receptor grid was generated using ASL residues of the top-ranked binding pocket. Peptides used as an inhibitor against Aβ42 are listed in Table 2. MM-GBSA method exhibit the free binding energy (ΔG bind) of each peptide inhibitor and is calculated using the following equation[38]:

**Table 1:**
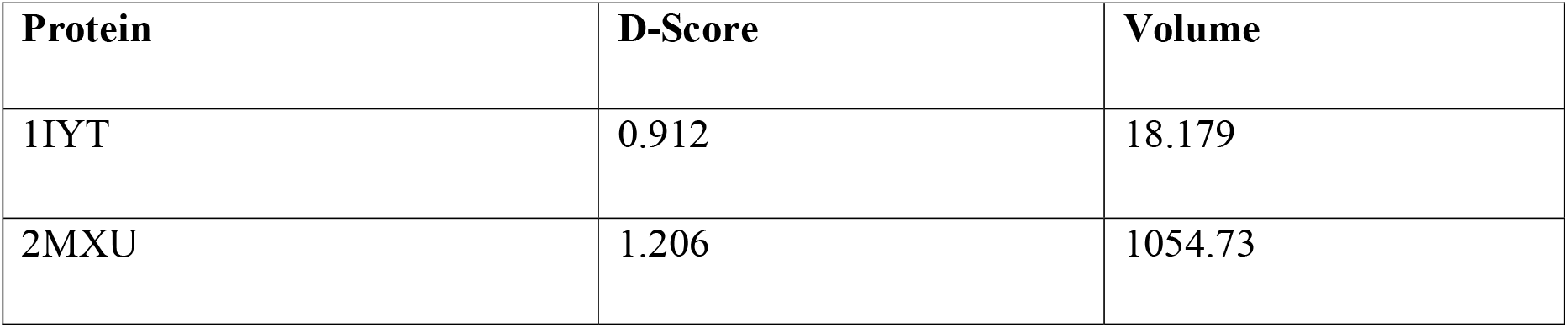
Binding Pocket score by SiteMap tool of Schroindger Suite

| Protein | D-Score | Volume |
| --- | --- | --- |
| 1IYT | 0.912 | 18.179 |
| 2MXU | 1.206 | 1054.73 |

**Table 2:**
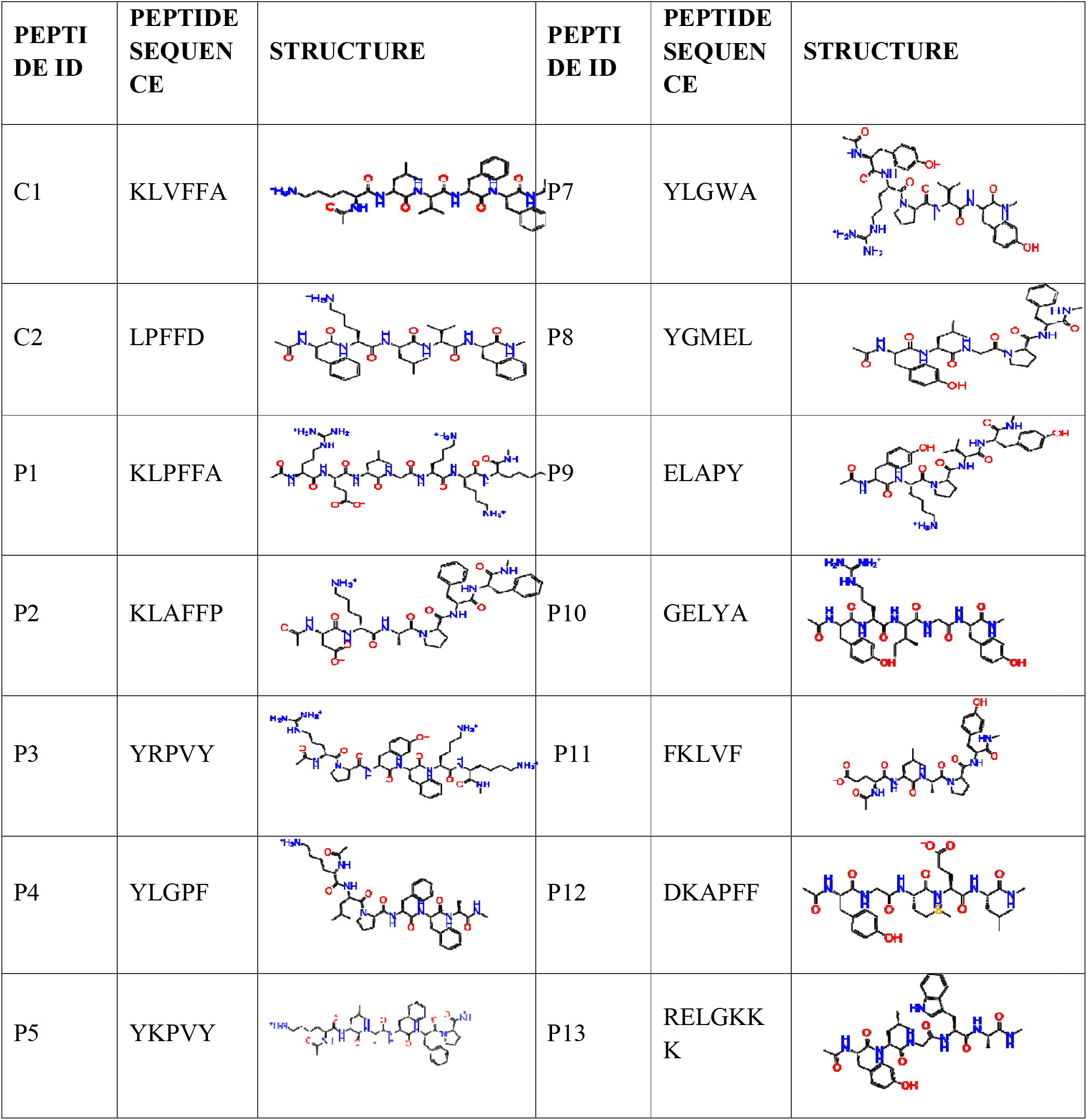
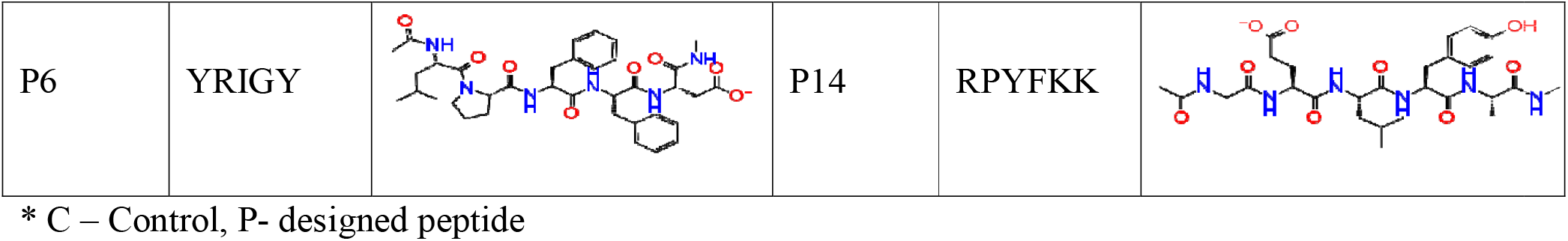
List of the designed peptides used for the study

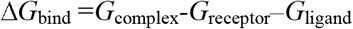

Higher the negative binding energy, stronger will be the ligand binding, and slower will be the oligomerization process. This is demonstrated by *Viet et al*. by determining the relationship between binding affinity and aggregation rate[22].

### 2. Molecular Dynamic Simulation

MD simulation is a powerful tool for studying the physicochemical properties of the biological system in terms of micro-degree of freedom [39]. We have studied the changes in structural stability upon the interaction between designed peptide-inhibitors and target proteins by performing classical MD simulations. As mentioned in table 4, P6 and P12 have a maximum binding affinity and were tested against the target proteins (Aβ42 monomer and fibril) using MD simulation. GROMACS [40]version 5.1.5 was used to carry out simulations using the charmm36-jul2020 force field, and the SPC water model was chosen for the system salvation [22]. The details of the simulations study have been enlisted in Table 3. The simulations of the target protein with inhibitors were performed in isobaric-isothermal assembly using periodic conditions. All 6 simulations were carried for 100ns and maintained the temperature at 300K and pressure at 1bar. Leapfrog algorithm was used for integrating the equation of motion with an integration time of 2fs. An identical simulation strategy was used for all the runs. It begins with relaxing the system in a vacuum via steepest descent minimization. Then water molecules (SOL) and ions for neutralizing the system were added and were relaxed by NVT MD for 1ps followed by NPT MD for the same period of 1ps. The trajectory analysis of the MD run was carried out by standard GROMACS command-based tools.

**Table 3:** Simulations Detail

| Simulation System | Simulation Time (ns) | no. of Atoms in protein | no. of Atoms in inhibitor | No. of atoms in complex | No. of SOL atoms | Na (sodium) Ions added | Total no. of atoms in a system |
| --- | --- | --- | --- | --- | --- | --- | --- |
| 1IYT (apoprotein) | 100 | 627 | - | - | 8934 x 3 | 3 | 27432 |
| 1IYT+ P6 |  | 627 | 104 | 731 | 8916 x 3 | 2 | 27481 |
| 1IYT+P12 |  | 627 | 110 | 737 | 8897 x 3 | 3 | 27431 |
| 2MXU (apoprotein) |  | 5736 | - | - | 30831 x 3 | 12 | 98241 |
| 2MXU+P6 |  | 5736 | 104 | 5840 | 22876 x 3 | 11 | 74479 |
| 2MXU+P12 |  | 5736 | 110 | 5846 | 22582 x 3 | 12 | 74414 |

**TABLE 4:** MM-GBSA Docking Result of designed peptide inhibitors to Aβ42 monomer(1IYT) and fibril (2MXU)

| Peptide ID | 2MXU (FIBRIL) |  | 1IYT (MONOMER) |  |
| --- | --- | --- | --- | --- |
|  | MM-GBSA score (kcal/mol) | Interacting Residues (aa\|residue no.\|chain no) | MM-GBSA score (kcal/mol) | Interacting Residues (aa\|residue no.) |
| C1 | -94.65 | His14F, Gly33D | -59.33 | Gly25, Glu22, **Ala21**, **Phe20** |
| C2 | -81.37 | His14C, His14E | -43.03 | Lys28, Glu22, **Ala21**, Ser26 |
| P2 | -106.02 | His14F, Gly33D, His14C,<br>His14E | -54.31 | Lys28, **Ala21** |
| P3 | -103.52 | His14G, Gly33G, His14E,<br>His14F | -47.77 | Gly38, Ala42, Glu22 |
| P4 | -102.39 | His14C, Gly33D | -55.85 | **Ala21**, Ser26 |
| P5 | -100.49 | His13F, His14E | -46.14 | **Lys16**, Glu22, Gly25,<br>Ser26 |
| P6 | -121.74 | Gly33G, His14F, His14E | -46.17 | Lys28, Gly22 |
| P7 | -95.04 | His14E, Gly33D, His14F | -61.70 | Lys28, Ser26, **Ala21** |
| P8 | -84.3 | Gly33D, His14E | -44.42 | Glu22 |
| P9 | -75.94 | Gly33C | -60.98 | Lys28, Ser26, Glu22,<br>**Phe20** |
| P10 | -77.39 | Gly33C | -46.85 | **Ala21**, Glu22, Ser26,<br>Lys28 |
| P11 | -101.59 | Gly33C, His14F | -47.36 | Glu22, Lys28 |
| P12 | -101.81 | His14F, His14E, His13E | -70.17 | **Ala21**, Lys28, Gly38,<br>Ala42 |
| P13 | -93.77 | - | -48.50 | - |
| P14 | -115.82 | - | -62.45 | - |

\* Amino acids highlighted in red lie within the predicted amyloidogenic regions of Aβ42 [43], [44]

## RESULTS

AD is a fatal neurodegenerative disorder and is considered a serious issue for society and the future health system[41]. Therefore, lots of researches have been carried out to find a better and promising solution for the inhibition of Aβ aggregation. In the present study, control peptides were modified, and their structural properties were used to design novel peptide inhibitors having high affinity and inhibitor efficiency against amyloid-beta fibrillization. These modifications were done in the following ways: (1) substituting proline as beta-breaker [42] (2) Acetylation of N-terminal and N-methylation of C-terminal (Ac-XXX-Nme) (3) Addition of TYR and PHE for improving binding affinity (4) C-terminal chain of lysine improves penetration power. Overall, 14 peptides were designed and evaluated primarily by docking, and the best inhibitor was simulated in the GROMACS package

### Molecular Docking Analysis

Docking helps to design novel drugs by estimating the binding position and binding affinity of the ligand with receptor/target. This study evaluates the binding interaction sites and affinity between Amyloid-beta monomer and aggregates with designed peptides using Schrodinger Suite - Maestro BioLuminate, and MM-GBSA tool is used for docking. Figure 3 shows the binding between the designed peptide having the highest binding affinity (P12) and Aβ42 monomer, whereas Figure 4 shows the binding between the designed peptide having the highest binding affinity (P6) with fibrillar Aβ42. Other interactions of representative structures are shown in the supplementary material. For more clarification on binding sites and their respective binding energy, the docking results for the best conformation of peptides are summarized in Table 4. From the data, on comparing with control peptide inhibitors, P6 shows the highest binding energy of -112.74 kcal/mol with 2MXU whereas, in the case of 1IYT, P12 shows maximum binding energy of -70.17 kcal/mol. Both the peptides lie within the best binding pocket and interact with nearby residues by H-bonding, Hydrophobic interactions, and, 1[-stacking. The interacting residues are mentioned in the docking result table. In the case of 1IYT, P12 interacts with Ala21, Lys28, Gly38, and Ala42 with H-bonding and also makes hydrophobic interaction with Ala42 and Lys28 (given in figure3a&3b). *Joker et al*. stated that there are three structural properties of interaction that can destabilize the assembly of Amyloid-beta: 1) Electrostatic interaction between residues Asp23-Lys28 forms a hydrophilic region; 2) Glu22 helps in maintaining the stability between chains of fibrils and helps to remain intact, and 3) Residue in between Lys17-Ala21 & Ala30-Val42 from a hydrophobic region which can be the target region to mask and block oligomerization [45]. As shown in figure-4b, upon docking, peptide (P6) interacts with the backbone of Amyloid-beta fibril (2MXU) that lies within the binding pocket having the maximum D-score (as mentioned in table1). P6 is found interacting with the His14 (E and F chain) and Gly33 (G chain) by three H-bonds. The complexity of 2MXU is much higher than other model amyloid-beta fibrils such as 2BEG. So far, no research has been done on the 2MXU amyloid-beta fibril. It has been stated by *Kanchi et al*. that histidine 14 has a sidechain of imidazole which helps in interacting with the cell membrane’s negatively charged phosphatidylserine molecule. Inhibiting this interaction helps in preventing Alzheimer’s Disease. Also, in preformed fibril states, His6, His13 & His14 molecules play a significant role in mediating zinc-induced oligomerization of fibrils. C1 peptide interacted with N-terminus Phe3, His14, Asp7, and Val12 and reported that it significantly destabilizes the amyloid-beta aggregates[46]. From the output, P6 and P12 were further tested for stability and interaction by MD Simulations which provide biological medium computationally and are used as a validation tool in CADD.

**Figure 3:**
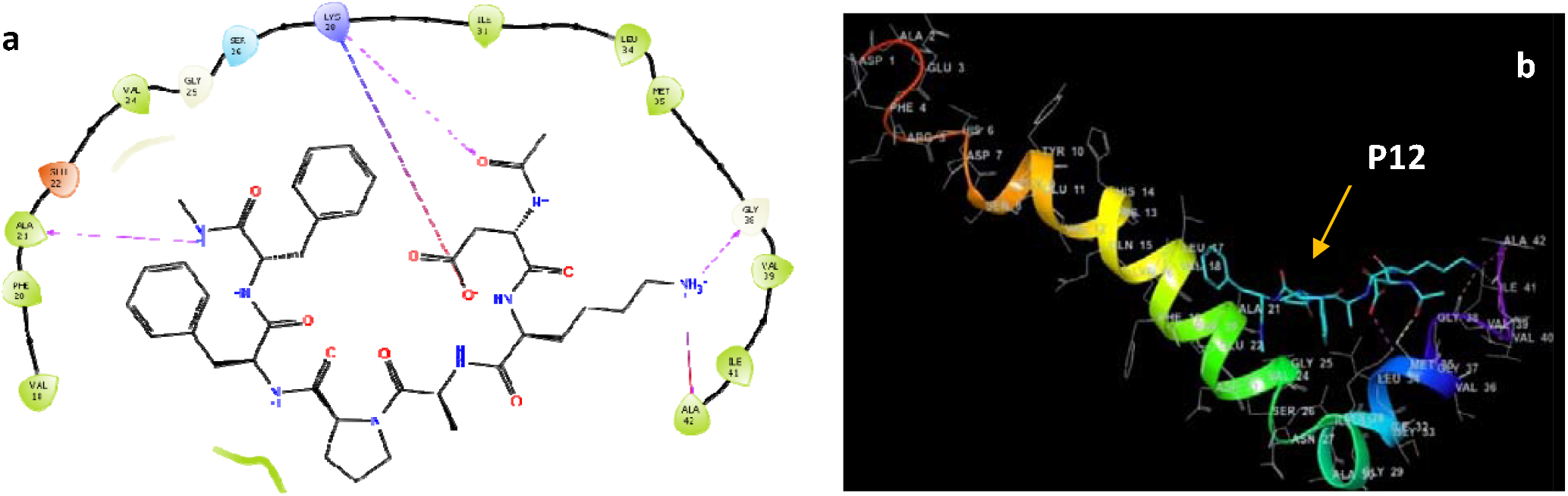
(a) 2D interaction layout and (b) 3D interaction between 1IYT and P12.

**Figure 4:**
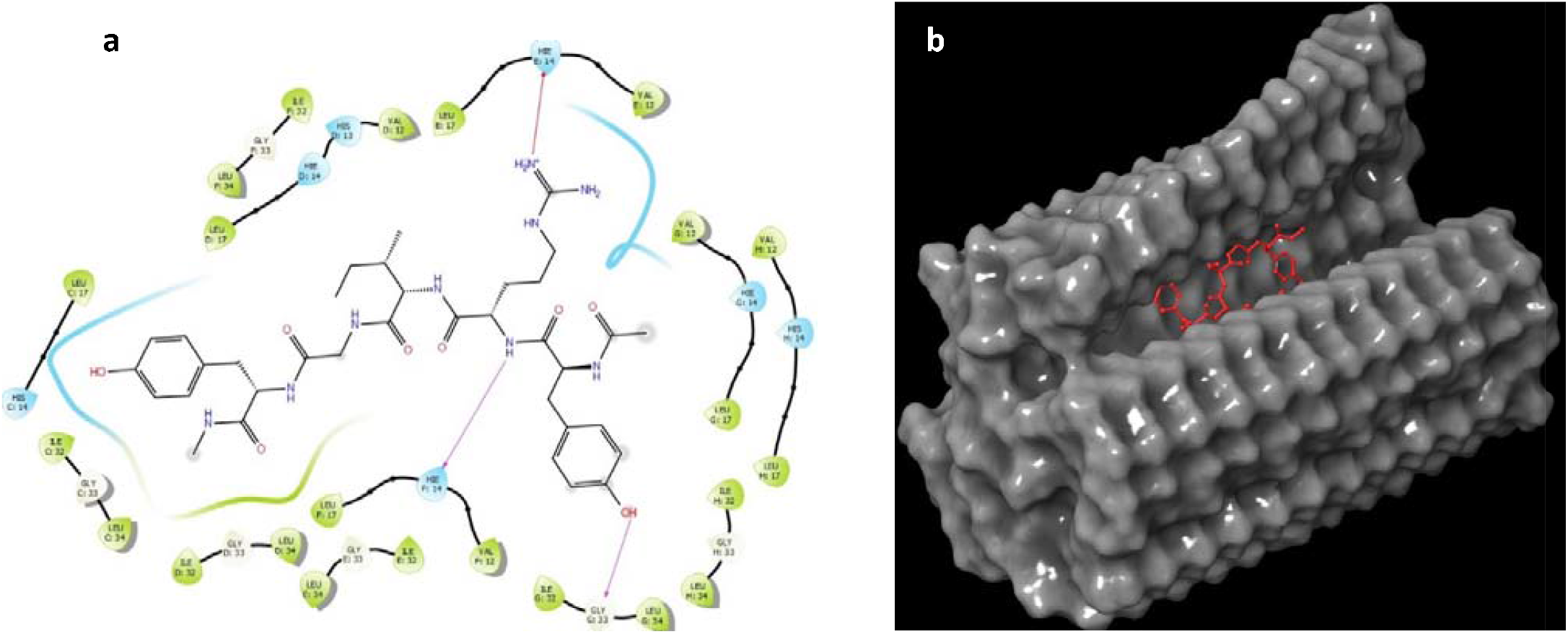
(a) 2D interaction layout and (b) surface 3D interaction between 2XMU and P6. P6 inhibitor interacts within binding pocket having D-score of 1.206.

### Molecular Dynamic Simulation Analysis

In this study, we performed 6-independent MD simulations on two different model proteins with the best-docked peptide to find the peptide having the highest potential for inhibiting the aggregation and disaggregation of fibrils. The conformational changes and respective stability of the system and involved protein were analyzed by four different parameters: RMSD, RMSF, Radius of Gyration, and H-bonding[47]. RMSD of backbone suggests the stability of protein and also predicts the conformational changes in protein within simulation time [48]. RMSF analysis helps to interpret the ratio of fluctuation at a residual level, and the Radius of gyration determines the compactness of protein. Steadier the values higher is its compactness, while values above reference point depict the unfolding/ loose packing of protein[49]. The protein which has a higher percentage of beta-sheets tends to form a stronger structure due to the presence of a large no. of H-bonds. Another MD analysis is based on the no. of H-bonds, in which if there is a decrease in H-bonds, then it represents that some kind of structural changes has occurred. Figure 5 shows the structural and the conformational changes of Aβ42 monomer (1IYT) with and without peptide inhibitors, and figure 6 shows changes in fibril (2MXU) with and without peptide inhibitors concerning simulation time. RMSD plot is shown in figure 7a; the monomeric peptide 1IYT (reference value) has an RMSD value of ∼0.7 nm at 25 ns, which is reduced to 0.55 nm during 42-50 ns, and again rises to 1.2 and get stabilized at 1.35 nm after 90 ns. In the presence of peptide inhibitor P6 at 30 ns, the sudden peak of ∼0.9-1 nm is observed, which shows destabilization, and as compared to the reference, it get stabilized after 95 ns. In the presence of peptide inhibitor P12, as compared with reference values, the peptide is initially stabilized at ∼0.3-0.4 nm, and from 13 ns onwards, some structural changes occurred as RMSD spiked to 0.7 nm. After 78 ns, there are no changes in structural conformation, and thus it gets stabilized. In the case of apoprotein 2MXU (figure 7b), the RMSD value is stabilized at ∼0.5 nm after 32 ns, while in the presence of P6 and P12, the values increased to ∼2.6-2.75 nm within a few ns, which indicated high destabilization/ distortion of the fibril. RMSF plot is given in figure 8; RMSF value in figure 8a is in the range of 0.2-0.9 nm, which shows the change in conformations on residues, whereas, in 1IYT alone, all the residues tend to change their conformations with time. In the presence of peptide inhibitors, RMSF shows different fluctuations compared to the reference values; P6 shows a fluctuation in the kink region at a greater extent, whereas in the presence of P12, it shows lesser fluctuation, which corresponds to higher stability. RMSF observation of fibril structure (figure 8b) indicates that in the presence of peptide inhibitor P6, structural destabilization occurs with great potential. The RMSF values of chains E, F, and G (Residues 170-220) spiked to 0.1nm to 0.38nm and 0.31nm, respectively. Fluctuations, to a greater extent, suggest distortion in fibrillar structure from the reported chains. Furthermore, the Radius of Gyration (Rg) was analyzed, which represents the compactness of protein. Higher the Rg value, less compact is the packing, and vice-versa[50]. Rg value below the reference point indicates tight packing or high compactness, whereas values above the reference point suggest the loose packing of proteins. As shown in figure 9a, the Rg value of 1IYT (apoprotein) is around 2 nm, whereas, in the presence of peptide inhibitors P6 & P12, the protein’s compactness increases as the value drops to 1.2 nm and 1.09 nm, respectively. In the case of fibril (2MXU) (Figure (b), the trend is opposite; in the presence of P6 & P12, the protein loses its compactness and becomes loosely packed. P6 shows more potential in converting the highly tight-packed fibril into a loosely packed distorted structure than P12. As reported earlier, proteins having beta-stands generally have a higher number of H-bonds than other secondary structures. Fibrils are highly stable beta-strand-rich molecules, so the decrease in the number of H-bonds suggests the structural transformation from beta-sheet to other secondary structures. Interaction between protein and ligand, protein-peptide in this study, is a must for both stabilization and destabilization, and the number of H-bonds after 100ns simulation is mentioned in table 5. The structure of 1IYT has 27 intramolecular H-bonds. In the presence of P6 lowers the number of H-bonds is seen as compared to P12. This is because of the presence of more H-bonds between P6 and the target protein. In the other case, 2MXU has 333 intramolecular H-bonds, and the number decreases in the presence of P6 and P12 to 272 and 283, respectively. From this, we can suggest that peptide inhibitors induce some structural changes to target proteins. The energy of interaction between protein and peptide inhibitor is reflected in the form of Coul-SR, and LJ-SR named electrostatic interaction energy and van der Waals force, respectively [51]. LJ-SR indicates non-bonded interaction energy. From table 6, it is observed that in both the cases (1IYT & 2MXU), P6 peptide inhibitor binding strongly with a target protein. Both Electrostatic and Vander Waal energies are higher when target protein bounded with P6 as compared with P12.

**Table 5:** H-bond Analysis

| Protein | Average no. of H-Bonds between<br>a target protein and peptide<br>inhibitor | Average no. of Intra H-bonds<br>after 50ns simulation |
| --- | --- | --- |
| 1IYT | - | 27.271 |
| 1IYT + P6 | 3.6 | 20.68 |
| 1IYT + P12 | 1.97 | 25.67 |
| 2MXU | - | 333.446 |
| 2MXU + P6 | 6.4 | 272.656 |
| 2MXU + P12 | 1.85 | 283.219 |

**6:** Interaction Energies

6: Interaction Energies
| PROTEIN+INHIBITOR | Coul-SR (kcal/mol) | LJ-SR (kcal/mol) |
| --- | --- | --- |
| 1IYT + P6 | -201.58 | -103.56 |
| 1IYT + P12 | -71.42 | -176.52 |
| 2MXU + P6 | -304.11 | -277.08 |
| 2MXU + P12 | -67.70 | -132.39 |

**Figure 5:**
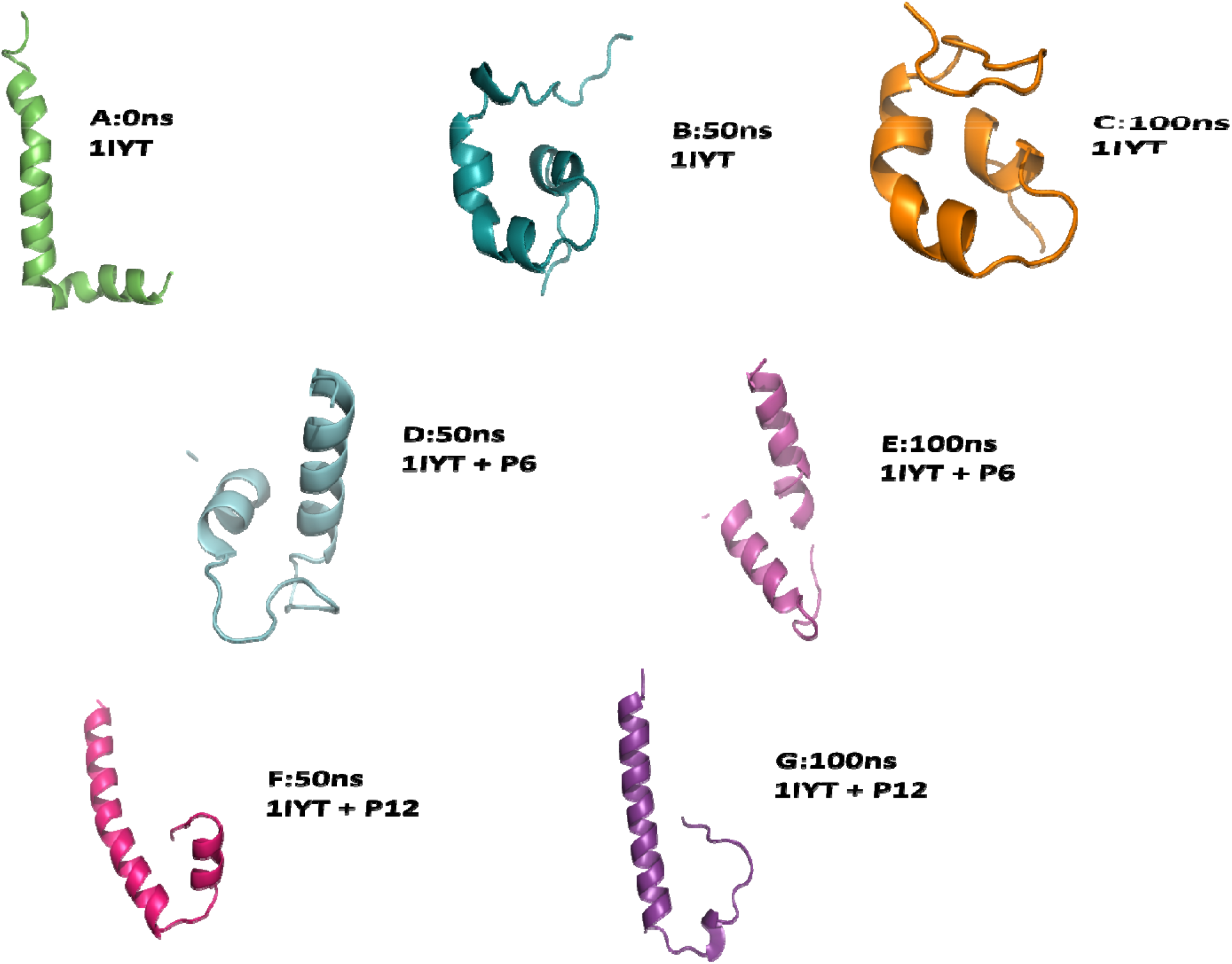
Structures of the production phase of MD simulation for 1IYT with and without peptide inhibitors. A depict two helical regions, after 50ns simulation as shown in B it gets converted into three helices connected with random coils and at 100ns 1IYT shows the highly complexed structure as shown in C. In the presence of P6 inhibitor initially, it gets converted into two helices as in D and then transformed into more stabilized helices connected with the coil having high helical content than protein structure at 50ns as in E. F and G are the representative structure of 1IYT in the presence of P12 which shows maintained helical integrity better than P6.

**Figure 6:**
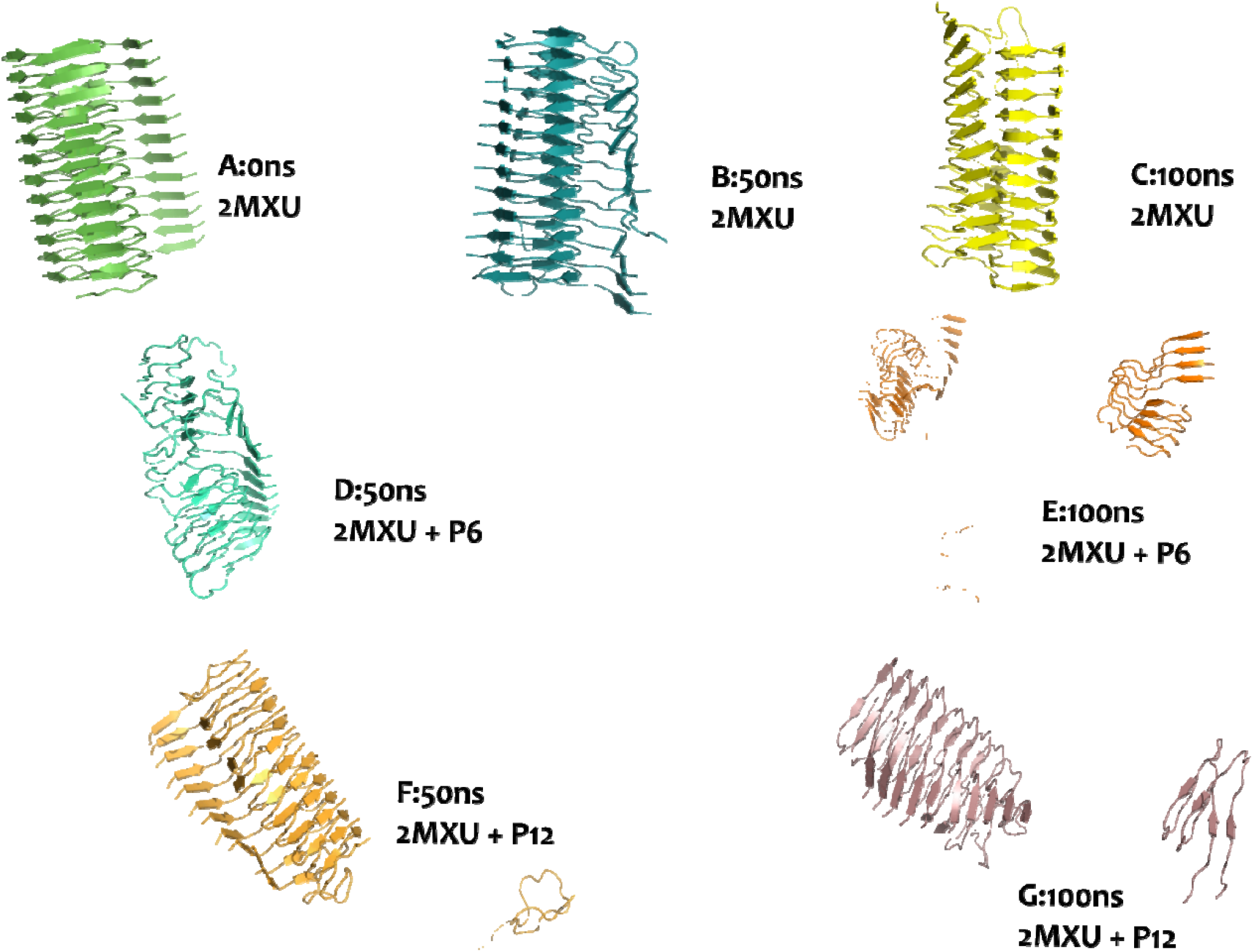
Structures of the production phase of MD simulation for 2MXU with and without peptide inhibitor. Without interacting with a peptide inhibitor, no conformational changes have been observed at 50ns and 100ns as shown in B & C in accordance to reference structure A. In the presence of P6, it shows structural distortion with the conversion of β-sheets converted into coils at 50ns as in D and at 100ns chains get separated from intact fibril structure as in E. Whereas with P12, at 50ns single-chain get separated as shown in F and two chains of intact 2MXU get separated at 100ns as in G. Thus, P6 has a higher potential for destabilizing the fibril structure than P12.

**Figure 7:**
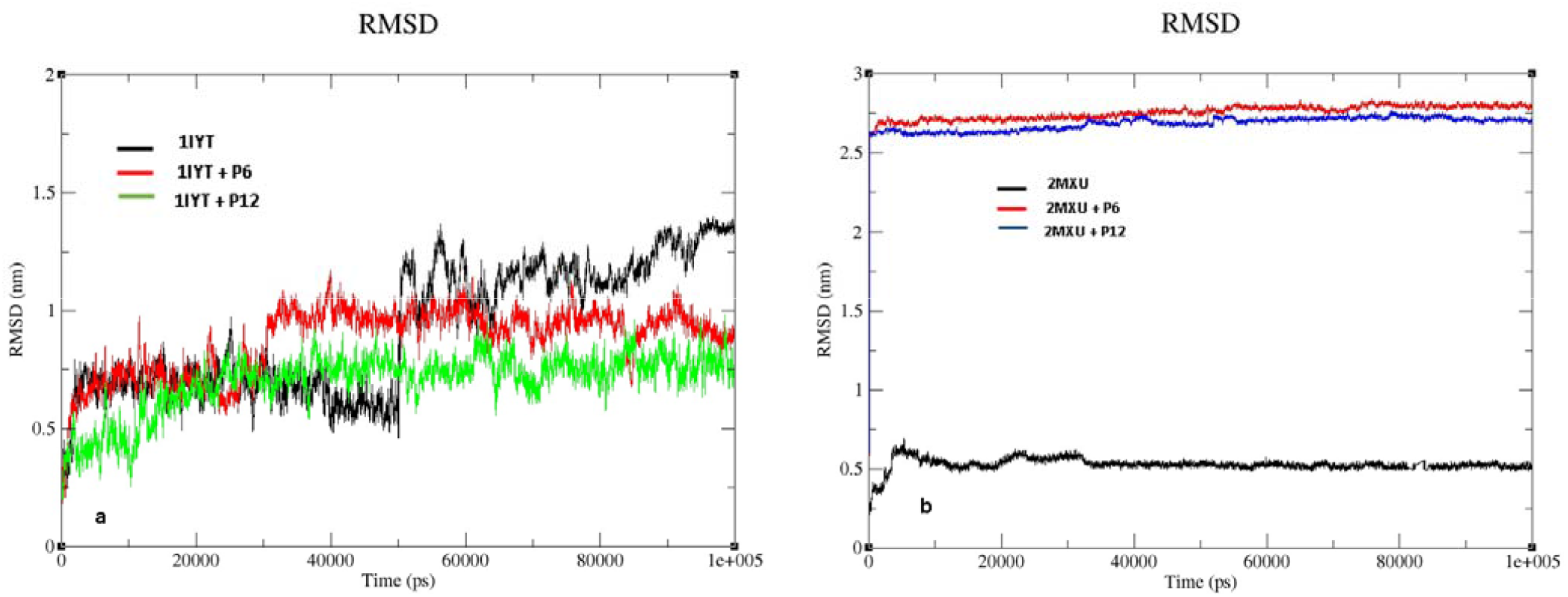
RMSD plot of 1IYT (a) and 2MXU (b) with and without the peptides

**Figure 8:**
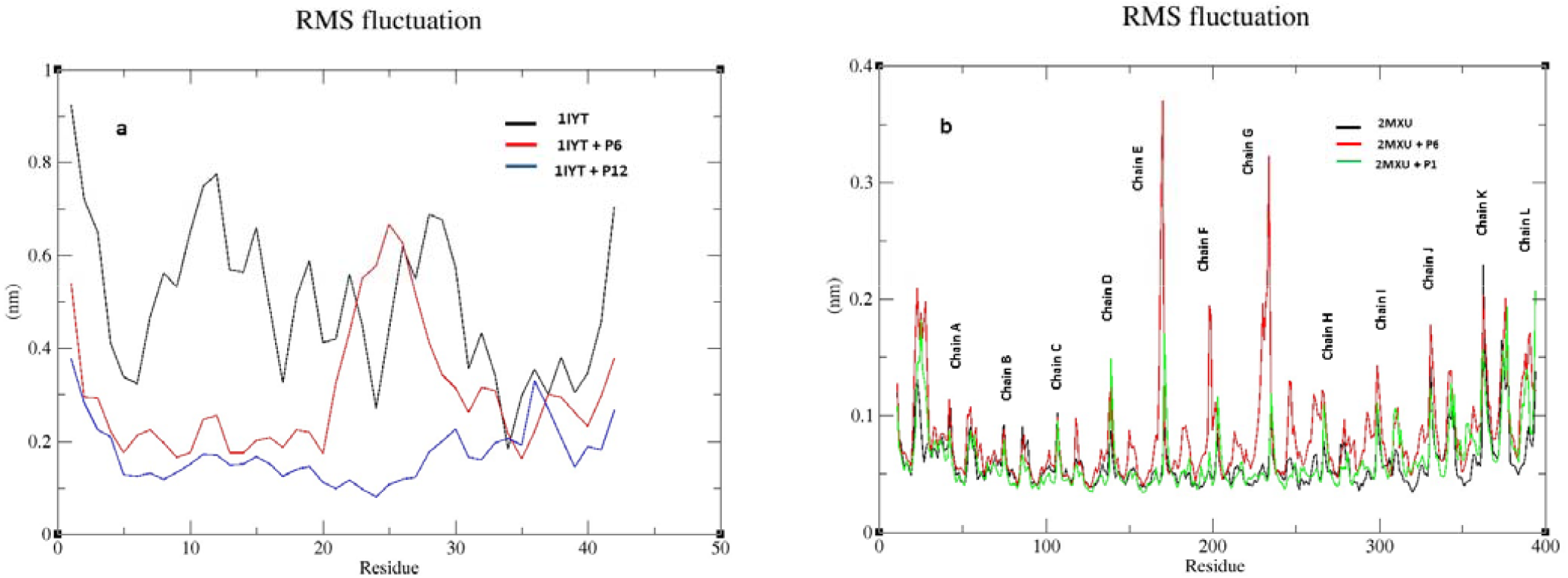
RMSF plot of 1IYT (A) and 2MXU (B) with and without the peptides

**Figure 9:**
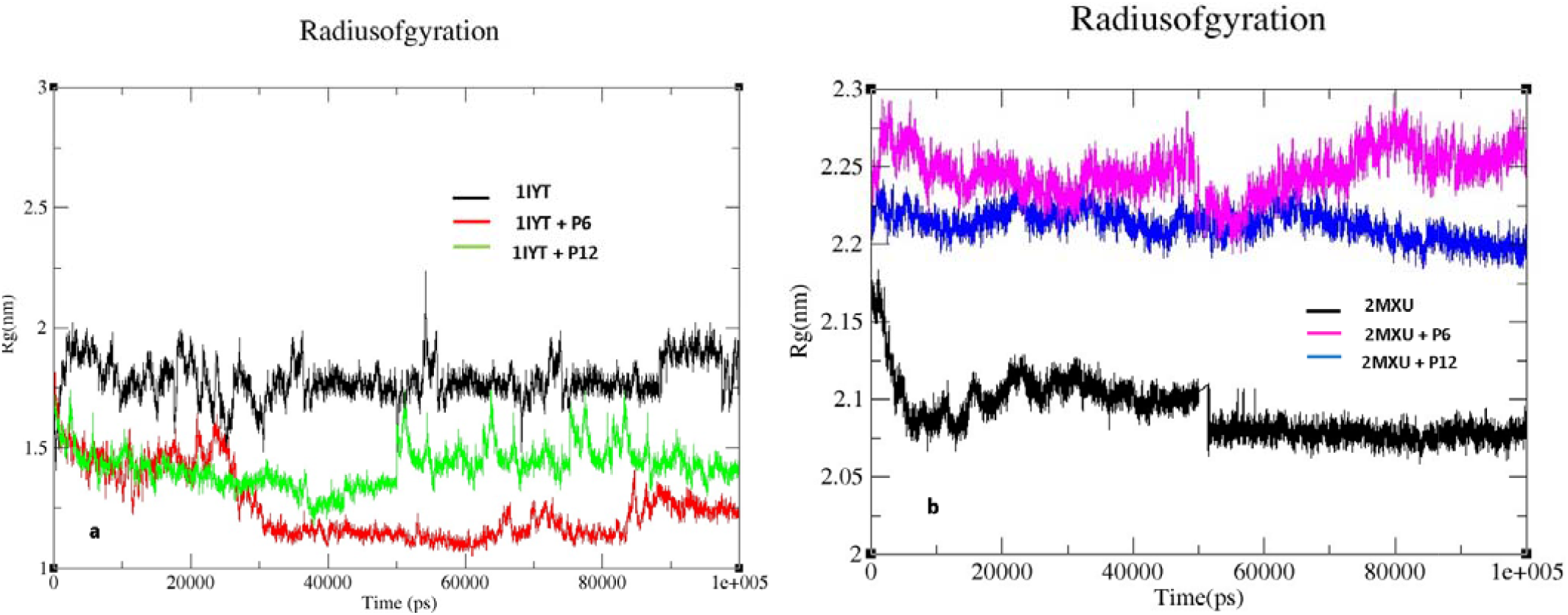
Radius of Gyration plot of 1IYT (A) and 2MXU (B) with and without the peptides

## DISCUSSION

Molecular docking is an essential tool for identifying novel therapeutics against amyloid-Beta fibrils leading to the prevention of AD. In combination with MD Simulations, the interaction between the novel designed peptides against Aβ42 fibrillization was studied. In this study, we compared the binding and structural stability of peptide inhibitors with both Aβ42 monomer and fibrils. For the first time, fibril structure (12 Chains) was used for estimation of interaction sites, binding affinity, structural stability in the presence/absence of peptide inhibitors. The MM-GBSA docking method showed that P6 has a higher binding affinity with Fibril (2MXU), whereas P12 has a higher affinity to Monomer (1IYT) of Aβ42. Structural stability analysis was done by performing six simulations for 100 ns. It was found that different conformations of P12 interact with monomer by binding with Ala21, Lys16, Lys28, Glu22, and ser26 residues. Interacting with these residues increases the stability of the native structure and is also responsible for blocking the nucleation sites by interacting particularly with Lys16 and Ala21 (residues lie within amyloidogenic regions) and thus inhibits the progression of oligomers and amyloid formation. Interaction energy calculated using *gmx* shows that P6 binds more strongly with both the target proteins than P12. In the case of the fibril, different conformations of P6 interact with Lys28, Asp23, Gly33, and His14. Binding with these residues helps in disrupting the highly stabilized fibrillar axis breaking the salt bridges. The mechanism behind both the steps, inhibition of amyloid formation and disaggregation of preformed fibrils, are quite different. By capping the regions which are responsible for the initiation of aggregation helps in blocking the sites leading to the prevent amyloid formation. In the case of stable fibrils, disaggregation and destabilization can be done by interaction with residues that lie within the fibrillar axis, breaking the salt bridges in between chains, and by interacting with residues in glycine grooves, etc. Therefore, from MD analysis by performing RMSD, RMSF, Rg, and H-bonds calculations, it was found that two different peptides have the potential for the modulation of amyloidogenesis. P12 has the potency to effectively inhibit the amyloid formation, and P6 has the potential for disaggregation of Amyloid-beta fibrils and can be considered as the best inhibitory peptide against Amyloid-Beta aggregation amongst the other peptides in our study. This work may provide computational insight into monomer stabilization and fibril disrupting mechanisms. From the gained information, in-vitro and in-vivo studies would be advantageous for identifying the effectiveness of peptides as a therapeutic agent against AD.

## ACKNOWLEDGEMENT

The authors acknowledge the infrastructural facility provided by NIT Rourkela. Further, the help of Mr. Praveen Kumar Guttala in the docking and simulations studies is acknowledged.

## REFERENCES

[1] M. Sunde and C. C. Blake, “From the globular to the fibrous state: protein structure and structural conversion in amyloid formation.,” Q. Rev. Biophys., vol. 31, no. 1, pp. 1–39, Feb. 1998, doi: 10.1017/s0033583598003400.

[2] C. M. Dobson, “Protein folding and misfolding.,” Nature, vol. 426, no. 6968, pp. 884–890, Dec. 2003, doi: 10.1038/nature02261.

[3] D. J. Selkoe, “The molecular pathology of Alzheimer’s disease,” Neuron, vol. 6, no. 4, pp. 487– 498, 1991, doi: https://doi.org/10.1016/0896-6273(91)90052-2.

[4] G. F. Chen et al., “Amyloid beta: Structure, biology and structure-based therapeutic development,” Acta Pharmacol. Sin., vol. 38, no. 9, pp. 1205–1235, 2017, doi: 10.1038/aps.2017.28.

[5] S. Weggen and D. Beher, “Molecular consequences of amyloid precursor protein and presenilin mutations causing autosomal-dominant Alzheimer’s disease.,” Alzheimers. Res. Ther., vol. 4, no. 2, p. 9, Mar. 2012, doi: 10.1186/alzrt107.

[6] J. A. Hardy and G. A. Higgins, “Alzheimer’s disease: the amyloid cascade hypothesis.,” Science, vol. 256, no. 5054, pp. 184–185, Apr. 1992, doi: 10.1126/science.1566067.

[7] C. Nilsberth et al., “The ‘Arctic’ APP mutation (E693G) causes Alzheimer’s disease by enhanced Abeta protofibril formation.,” Nat. Neurosci., vol. 4, no. 9, pp. 887–893, Sep. 2001, doi: 10.1038/nn0901-887.

[8] L. Gu and Z. Guo, “Alzheimer’s Aβ42 and Aβ40 peptides form interlaced amyloid fibrils.,” J. Neurochem., vol. 126, no. 3, pp. 305–311, Aug. 2013, doi: 10.1111/jnc.12202.

[9] I. Morgado and M. Garvey, “Lipids in Amyloid-β Processing, Aggregation, and Toxicity.,” Adv. Exp. Med. Biol., vol. 855, pp. 67–94, 2015, doi: 10.1007/978-3-319-17344-3_3.

[10] C. A. Paredes-Rosan, D. E. Valencia, H. L. Barazorda-Ccahuana, J. A. Aguilar-Pineda, and B. Gómez, “Amyloid beta oligomers: how pH influences over trimer and pentamer structures?,” J. Mol. Model., vol. 26, no. 1, p. 1, Dec. 2019, doi: 10.1007/s00894-019-4247-5.

[11] S. P. Radko et al., “Heparin Modulates the Kinetics of Zinc-Induced Aggregation of Amyloid-β Peptides.,” J. Alzheimers. Dis., vol. 63, no. 2, pp. 539–550, 2018, doi: 10.3233/JAD-171120.

[12] Y. Bin, X. Li, Y. He, S. Chen, and J. Xiang, “Amyloid-β peptide (1-42) aggregation induced by copper ions under acidic conditions.,” Acta Biochim. Biophys. Sin. (Shanghai)., vol. 45, no. 7, pp. 570–577, Jul. 2013, doi: 10.1093/abbs/gmt044.

[13] J. Roche, Y. Shen, J. H. Lee, J. Ying, and A. Bax, “Monomeric Aβ(1-40) and Aβ(1-42) Peptides in Solution Adopt Very Similar Ramachandran Map Distributions That Closely Resemble Random Coil.,” Biochemistry, vol. 55, no. 5, pp. 762–775, Feb. 2016, doi: 10.1021/acs.biochem.5b01259.

[14] J. A. White, A. M. Manelli, K. H. Holmberg, L. J. Van Eldik, and M. J. Ladu, “Differential effects of oligomeric and fibrillar amyloid-beta 1-42 on astrocyte-mediated inflammation.,” Neurobiol. Dis., vol. 18, no. 3, pp. 459–465, Apr. 2005, doi: 10.1016/j.nbd.2004.12.013.

[15] D. V Hansen, J. E. Hanson, and M. Sheng, “Microglia in Alzheimer’s disease.,” J. Cell Biol., vol. 217, no. 2, pp. 459–472, Feb. 2018, doi: 10.1083/jcb.201709069.

[16] D. Giulian et al., “The HHQK domain of beta-amyloid provides a structural basis for the immunopathology of Alzheimer’s disease.,” J. Biol. Chem., vol. 273, no. 45, pp. 29719–29726, Nov. 1998, doi: 10.1074/jbc.273.45.29719.

[17] S. O. Garbuzynskiy, M. Y. Lobanov, and O. V Galzitskaya, “FoldAmyloid: a method of prediction of amyloidogenic regions from protein sequence.,” Bioinformatics, vol. 26, no. 3, pp. 326–332, Feb. 2010, doi: 10.1093/bioinformatics/btp691.

[18] N. S. de Groot, V. Castillo, R. Graña-Montes, and S. Ventura, “AGGRESCAN: method, application, and perspectives for drug design.,” Methods Mol. Biol., vol. 819, pp. 199–220, 2012, doi: 10.1007/978-1-61779-465-0_14.

[19] G. Eskici and M. Gur, “Computational Design of New Peptide Inhibitors for Amyloid Beta (Aβ) Aggregation in Alzheimer’s Disease: Application of a Novel Methodology,” PLoS One, vol. 8, no. 6, 2013, doi: 10.1371/journal.pone.0066178.

[20] K. Wiesehan, J. Stöhr, L. Nagel-Steger, T. van Groen, D. Riesner, and D. Willbold, “Inhibition of cytotoxicity and amyloid fibril formation by a D-amino acid peptide that specifically binds to Alzheimer’s disease amyloid peptide.,” Protein Eng. Des. Sel., vol. 21, no. 4, pp. 241–246, Apr. 2008, doi: 10.1093/protein/gzm054.

[21] T. Arai, D. Sasaki, T. Araya, T. Sato, Y. Sohma, and M. Kanai, “A cyclic KLVFF-derived peptide aggregation inhibitor induces the formation of less-toxic off-pathway amyloid-β oligomers.,” Chembiochem, vol. 15, no. 17, pp. 2577–2583, Nov. 2014, doi: 10.1002/cbic.201402430.

[22] M. H. Viet, S. T. Ngo, N. S. Lam, and M. S. Li, “Inhibition of aggregation of amyloid peptides by beta-sheet breaker peptides and their binding affinity,” J. Phys. Chem. B, vol. 115, no. 22, pp. 7433–7446, 2011, doi: 10.1021/jp1116728.

[23] C. Soto, E. M. Sigurdsson, L. Morelli, R. A. Kumar, E. M. Castaño, and B. Frangione, “Beta-sheet breaker peptides inhibit fibrillogenesis in a rat brain model of amyloidosis: implications for Alzheimer’s therapy.,” Nat. Med., vol. 4, no. 7, pp. 822–826, Jul. 1998, doi: 10.1038/nm0798-822.

[24] I. Granic et al., “LPYFDa neutralizes amyloid-beta-induced memory impairment and toxicity.,” J. Alzheimers. Dis., vol. 19, no. 3, pp. 991–1005, 2010, doi: 10.3233/JAD-2010-1297.

[25] C. Soto, “sheet breaker peptides dissolving the therapeutic problem of Alzheimer ‘ s disease[?,” 2002.

[26] A. Orjuela et al., “Computational Evaluation of Interaction Between Curcumin Derivatives and Amyloid-β Monomers and Fibrils: Relevance to Alzheimer’s Disease,” J. Alzheimer’s Dis., vol. Preprint, pp. 1–13, 2020, doi: 10.3233/JAD-200941.

[27] A. Mitra and N. Sarkar, “Sequence and structure-based peptides as potent amyloid inhibitors: A review,” Arch. Biochem. Biophys., vol. 695, p. 108614, 2020, doi: https://doi.org/10.1016/j.abb.2020.108614.

[28] J. Huang et al., “CHARMM36m: an improved force field for folded and intrinsically disordered proteins,” Nat. Methods, vol. 14, no. 1, pp. 71–73, 2017, doi: 10.1038/nmeth.4067.

[29] Y. Wu, H. L. Tepper, and G. A. Voth, “Flexible simple point-charge water model with improved liquid-state properties,” J. Chem. Phys., vol. 124, no. 2, 2006, doi: 10.1063/1.2136877.

[30] K. Zhu, T. Day, D. Warshaviak, C. Murrett, R. Friesner, and D. Pearlman, “Antibody structure determination using a combination of homology modeling, energy-based refinement, and loop prediction.,” Proteins, vol. 82, no. 8, pp. 1646–1655, Aug. 2014, doi: 10.1002/prot.24551.

[31] D. Seeliger and B. L. de Groot, “Ligand docking and binding site analysis with PyMOL and Autodock/Vina.,” J. Comput. Aided. Mol. Des., vol. 24, no. 5, pp. 417–422, May 2010, doi: 10.1007/s10822-010-9352-6.

[32] W. Humphrey, A. Dalke, and K. Schulten, “VMD: visual molecular dynamics.,” J. Mol. Graph., vol. 14, no. 1, pp. 27-28,33-38, Feb. 1996, doi: 10.1016/0263-7855(96)00018-5.

[33] D. Van Der Spoel, E. Lindahl, B. Hess, G. Groenhof, A. E. Mark, and H. J. C. Berendsen, “GROMACS: fast, flexible, and free.,” J. Comput. Chem., vol. 26, no. 16, pp. 1701–1718, Dec. 2005, doi: 10.1002/jcc.20291.

[34] K. Roy, S. Kar, and R. N. Das, “Chapter 10 -Other Related Techniques,” K. Roy, S. Kar, and R. N. B. T.-U. the B. of Q. for A. in P. S. and R. A. Das, Eds. Boston: Academic Press, 2015, pp. 357–425.

[35] H. Beard, A. Cholleti, D. Pearlman, W. Sherman, and K. A. Loving, “Applying physics-based scoring to calculate free energies of binding for single amino acid mutations in protein-protein complexes,” PLoS One, vol. 8, no. 12, pp. 1–11, 2013, doi: 10.1371/journal.pone.0082849.

[36] N. K. Salam, M. Adzhigirey, W. Sherman, D. A. Pearlman, and D. Thirumalai, “Structure-based approach to the prediction of disulfide bonds in proteins,” Protein Eng. Des. Sel., vol. 27, no. 10, pp. 365–374, 2014, doi: 10.1093/protein/gzu017.

[37] G. Sanchez, “Las instituciones de ciencia y tecnología en los procesos de aprendizaje de la producción agroalimentaria en Argentina,” El Sist. argentino innovación Inst. Empres. y redes. El desafío la creación y apropiación Conoc., 2013, doi: 10.1002/prot.

[38] S. V. Pattar, S. A. Adhoni, C. M. Kamanavalli, and S. S. Kumbar, “In silico molecular docking studies and MM/GBSA analysis of coumarin-carbonodithioate hybrid derivatives divulge the anticancer potential against breast cancer,” Beni-Suef Univ. J. Basic Appl. Sci., vol. 9, no. 1, 2020, doi: 10.1186/s43088-020-00059-7.

[39] V. Minicozzi et al., “Computational and Experimental Studies on [-Sheet Breakers Targeting A [1 – 40 Fibrils,” 2014, doi: 10.1074/jbc.M113.537472.

[40] M. J. Abraham et al., “GROMACS: High performance molecular simulations through multi-level parallelism from laptops to supercomputers,” SoftwareX, vol. 1–2, pp. 19–25, 2015, doi: https://doi.org/10.1016/j.softx.2015.06.001.

[41] J. R. Horsley, B. Jovcevski, K. L. Wegener, J. Yu, T. L. Pukala, and A. D. Abell, “Rationally designed peptide-based inhibitor of Aβ42fibril formation and toxicity: A potential therapeutic strategy for Alzheimer’s disease,” Biochem. J., vol. 477, no. 11, pp. 1541–1564, 2020, doi: 10.1042/BCJ20200290.

[42] J. F. Poduslo, G. L. Curran, A. Kumar, B. Frangione, and C. Soto, “Beta-sheet breaker peptide inhibitor of Alzheimer’s amyloidogenesis with increased blood-brain barrier permeability and resistance to proteolytic degradation in plasma.,” J. Neurobiol., vol. 39, no. 3, pp. 371–382, Jun. 1999.

[43] A. Henning-Knechtel et al., “Designed Cell-Penetrating Peptide Inhibitors of Amyloid-beta Aggregation and Cytotoxicity,” Cell Reports Phys. Sci., vol. 1, no. 2, p. 100014, 2020, doi: 10.1016/j.xcrp.2020.100014.

[44] B. Neddenriep et al., “Short Peptides as Inhibitors of Amyloid Aggregation.,” Open Biotechnol. J., vol. 5, pp. 39–46, Dec. 2011, doi: 10.2174/1874070701105010039.

[45] S. Jokar et al., “Design of peptide-based inhibitor agent against amyloid-β aggregation: Molecular docking, synthesis and in vitro evaluation.,” Bioorg. Chem., vol. 102, p. 104050, Sep. 2020, doi: 10.1016/j.bioorg.2020.104050.

[46] P. K. Kanchi and A. K. Dasmahapatra, “Enhancing the binding of the β-sheet breaker peptide LPFFD to the amyloid-β fibrils by aromatic modifications: A molecular dynamics simulation study,” Comput. Biol. Chem., vol. 92, no. October 2020, p. 107471, 2021, doi: 10.1016/j.compbiolchem.2021.107471.

[47] K. Al-Khafaji, D. AL-Duhaidahawi, and T. Taskin Tok, “Using integrated computational approaches to identify safe and rapid treatment for SARS-CoV-2,” J. Biomol. Struct. Dyn., vol. 39, no. 9, pp. 3387–3395, Jun. 2021, doi: 10.1080/07391102.2020.1764392.

[48] K. Sargsyan, C. Grauffel, and C. Lim, “How Molecular Size Impacts RMSD Applications in Molecular Dynamics Simulations.,” J. Chem. Theory Comput., vol. 13, no. 4, pp. 1518–1524, Apr. 2017, doi: 10.1021/acs.jctc.7b00028.

[49] Y. Zhao, C. Zeng, and M. A. Massiah, “Molecular Dynamics Simulation Reveals Insights into the Mechanism of Unfolding by the A130T/V Mutations within the MID1 Zinc-Binding Bbox1 Domain,” PLoS One, vol. 10, no. 4, p. e0124377, Apr. 2015, [Online]. Available: https://doi.org/10.1371/journal.pone.0124377.

[50] M. Y. Lobanov, N. S. Bogatyreva, and O. V Galzitskaya, “Radius of gyration as an indicator of protein structure compactness,” Mol. Biol., vol. 42, no. 4, pp. 623–628, 2008, doi: 10.1134/S0026893308040195.

[51] I. Doytchinova, P. Petkov, I. Dimitrov, M. Atanasova, and D. R. Flower, “HLA-DP2 binding prediction by molecular dynamics simulations.,” Protein Sci., vol. 20, no. 11, pp. 1918–1928, Nov. 2011, doi: 10.1002/pro.732.

