## Supplementary Material for "*In silico* identification of novel peptides as potential modulators of Aβ42 Amyloidogenesis"

1.
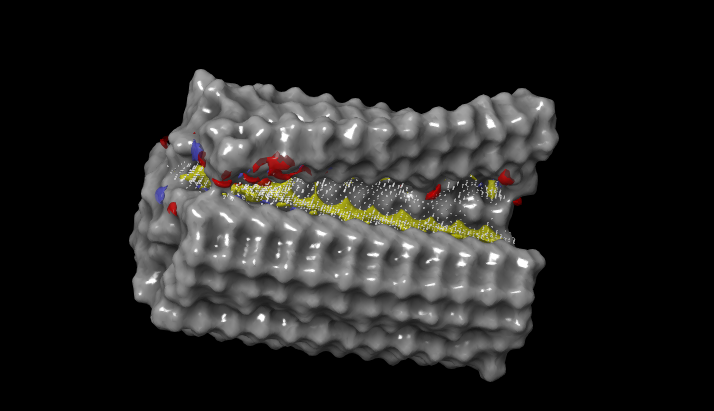
**Binding Pockets predicted by SiteMap tool of Schrodinge**
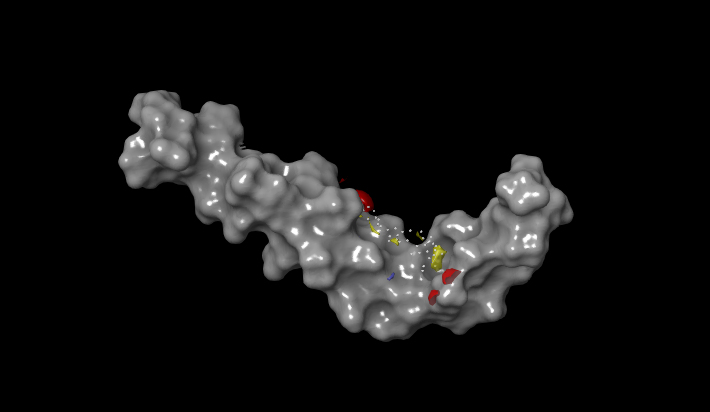
**r**

****Red/Yellow area represents binding pocket***

a) **2MXU**
b) **1IYT**
2. **2D Interaction images of 1IYT with designed peptide inhibitors P1**-**P13 (A**🡪 **M)**

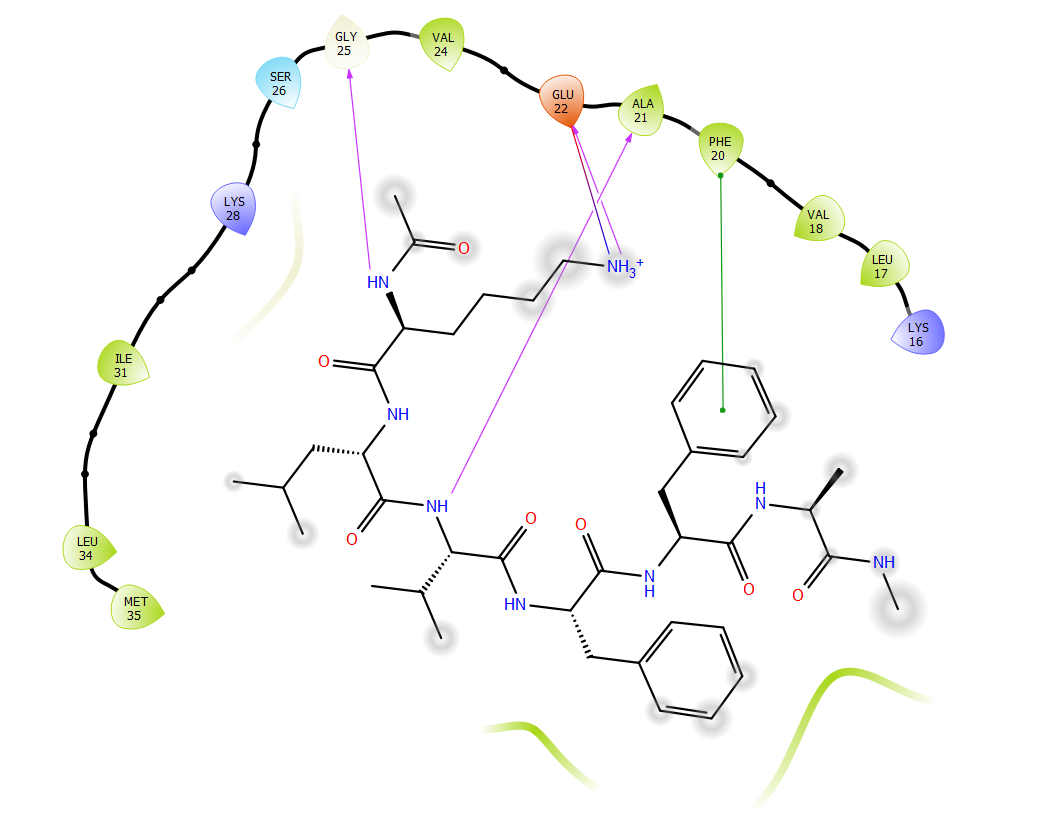

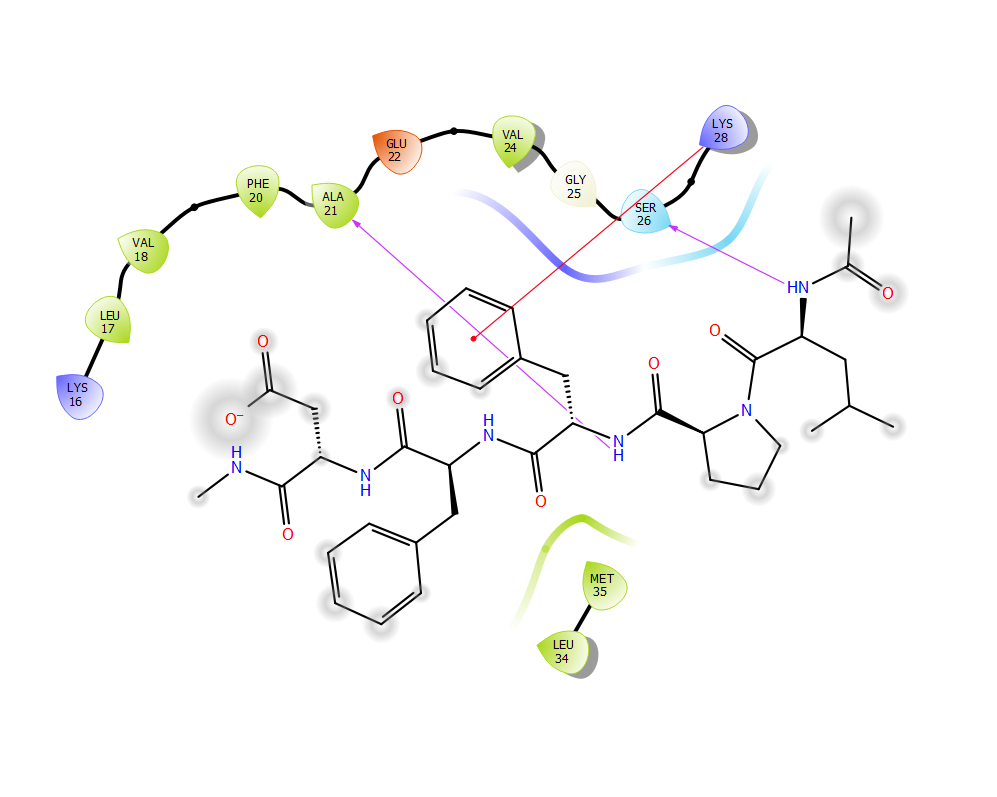

A) **C2**
B) **C1**

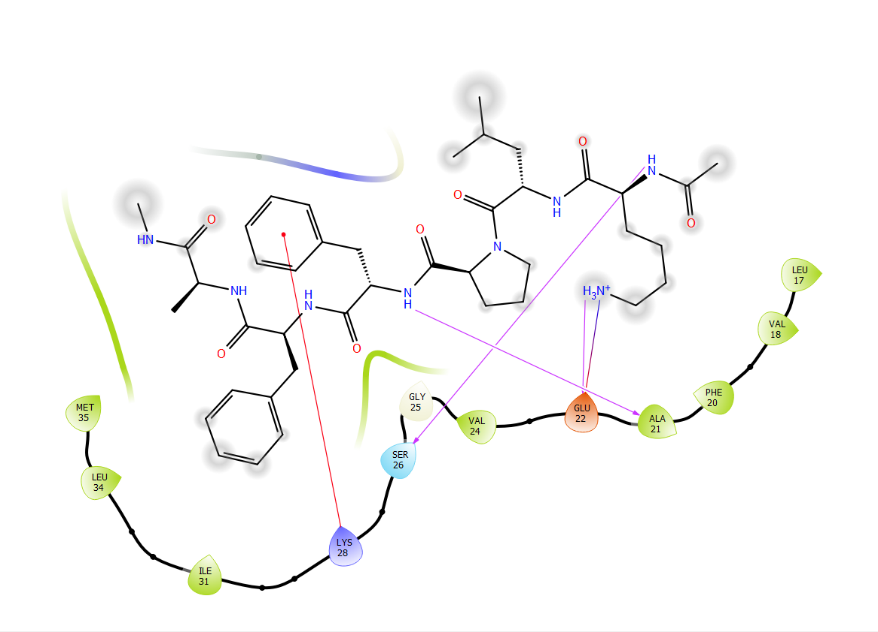

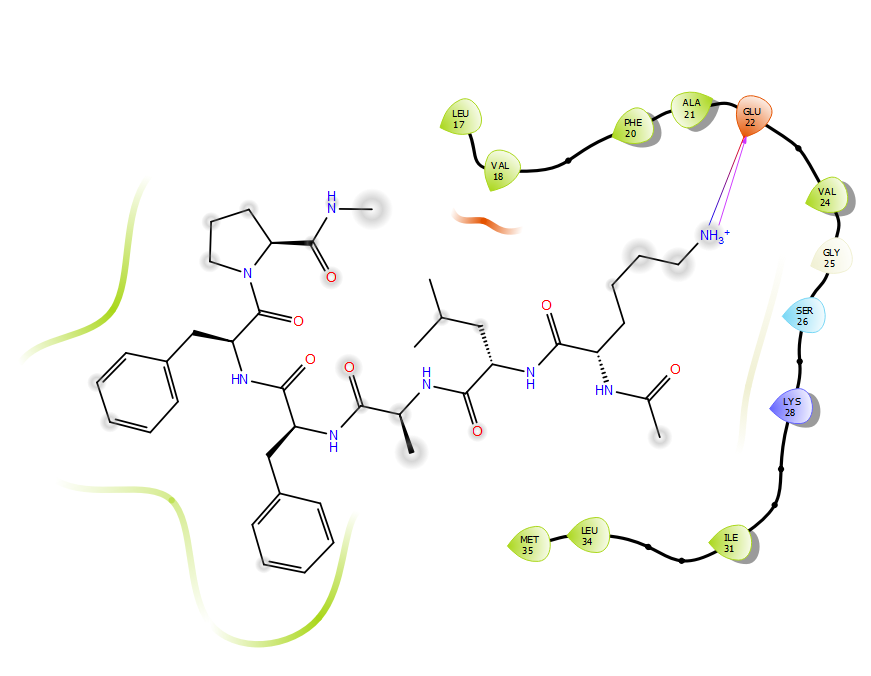

C) **P2**
D) **P1**

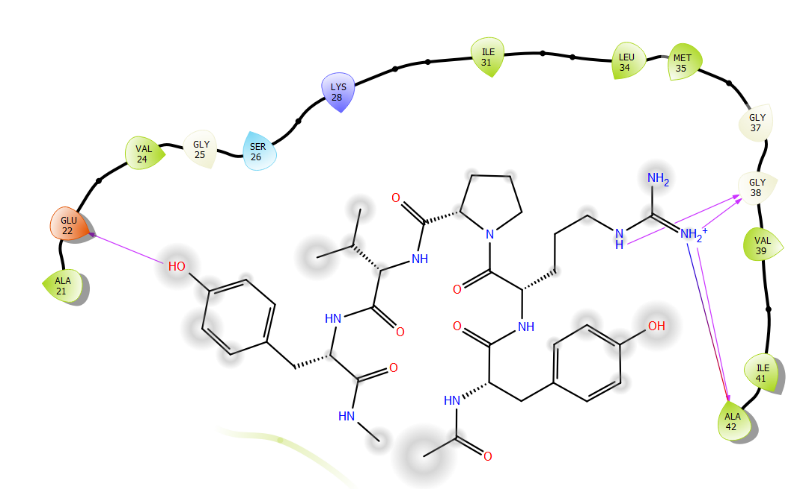

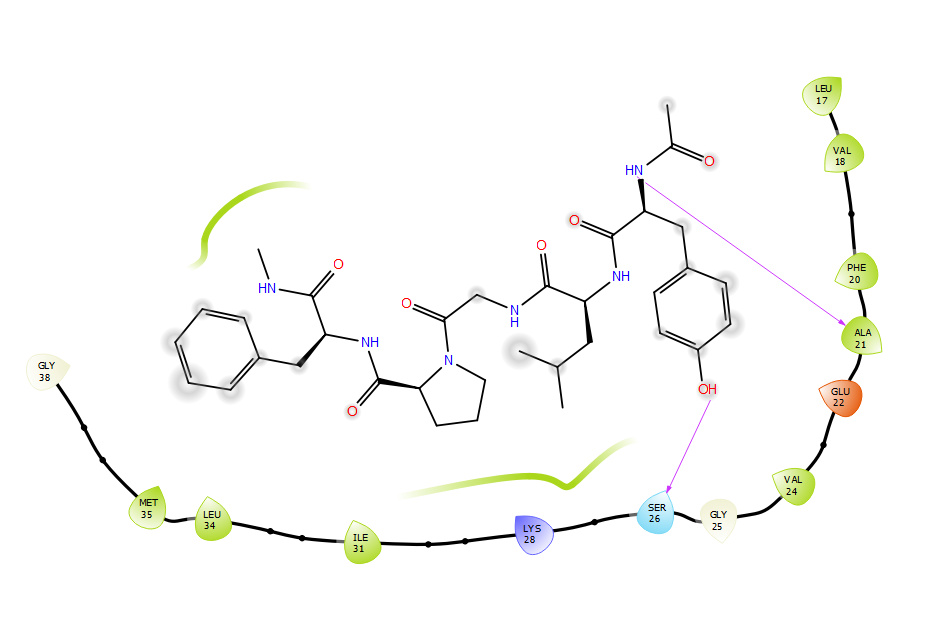

E) **P4**
F) **P3**

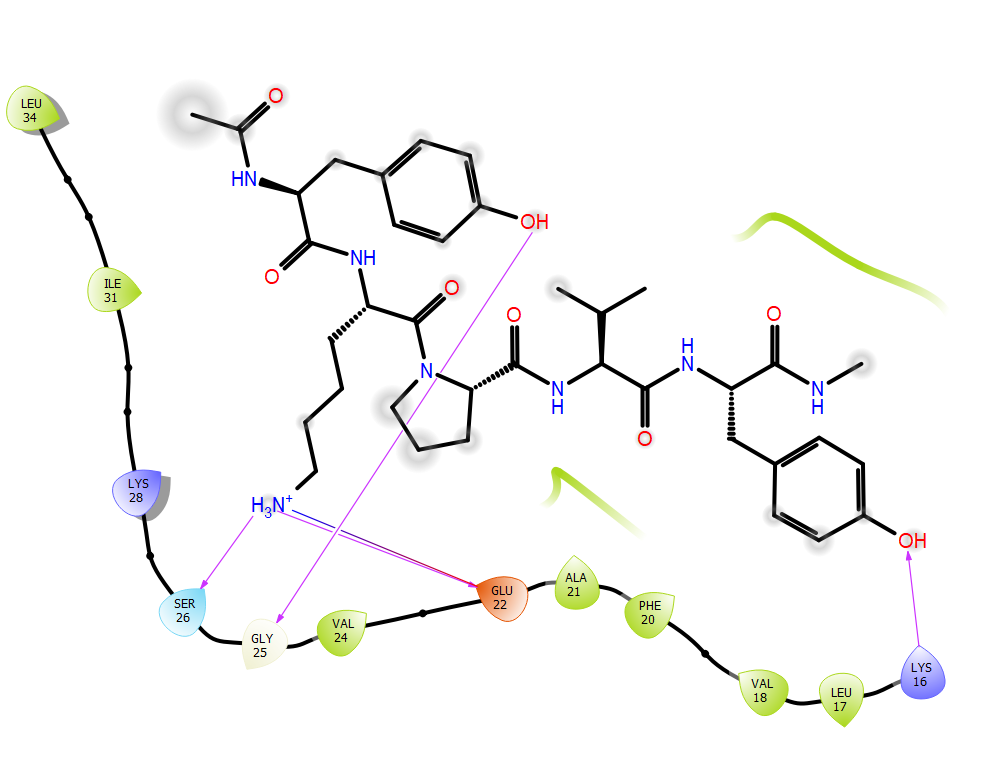

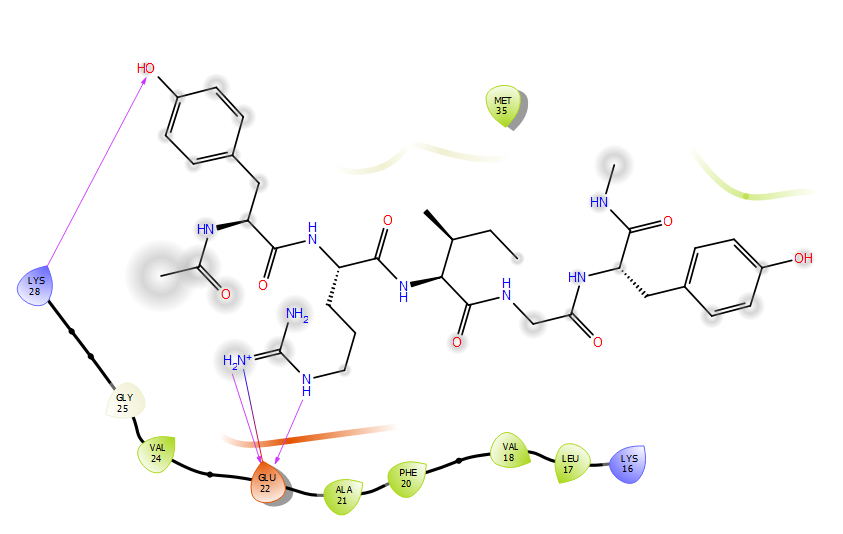

G) **P6**
H) **P5**

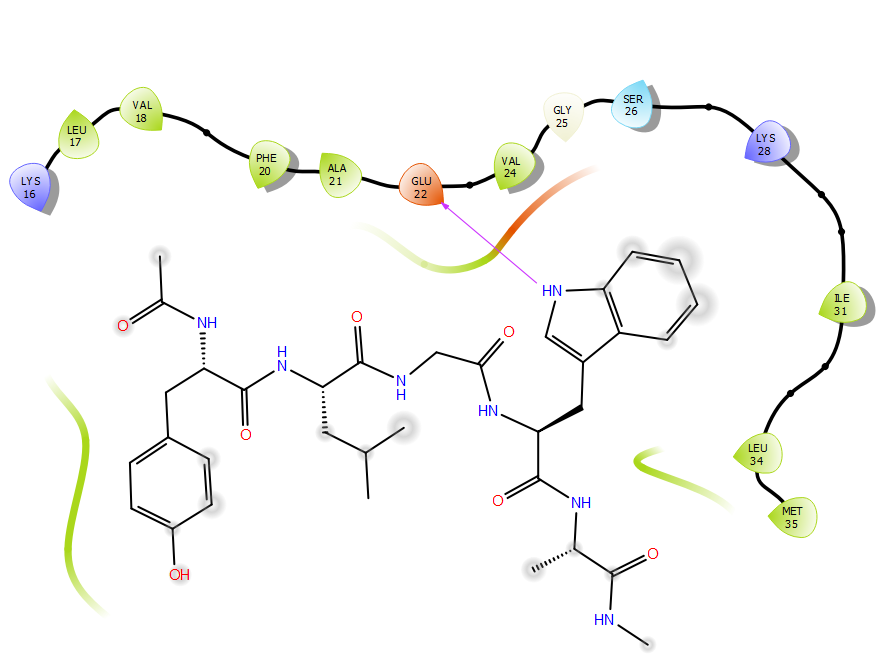

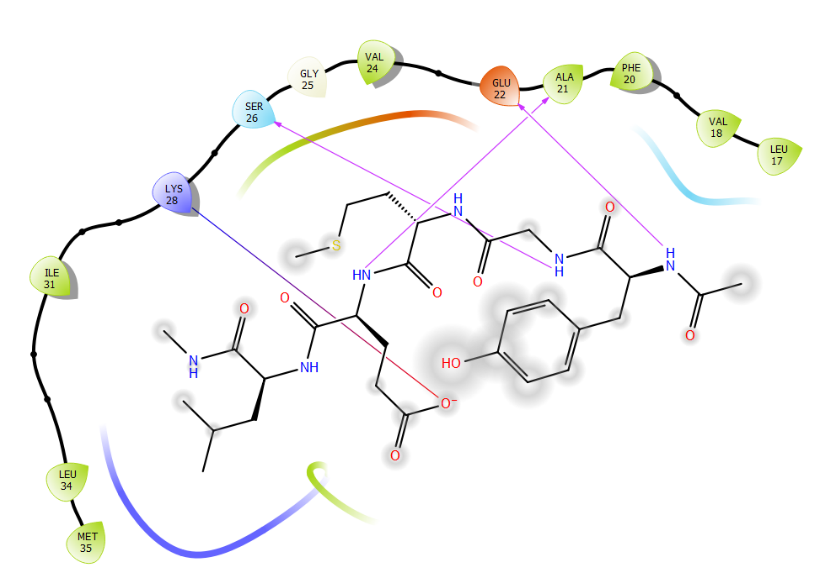

I) **P8**
J) **P7**

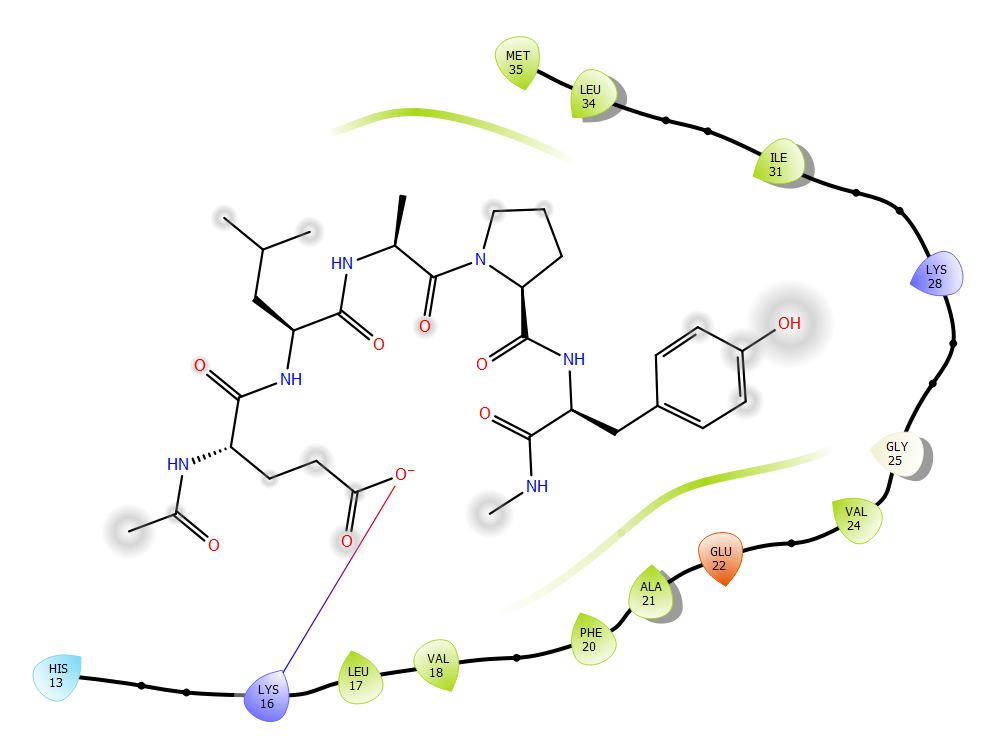

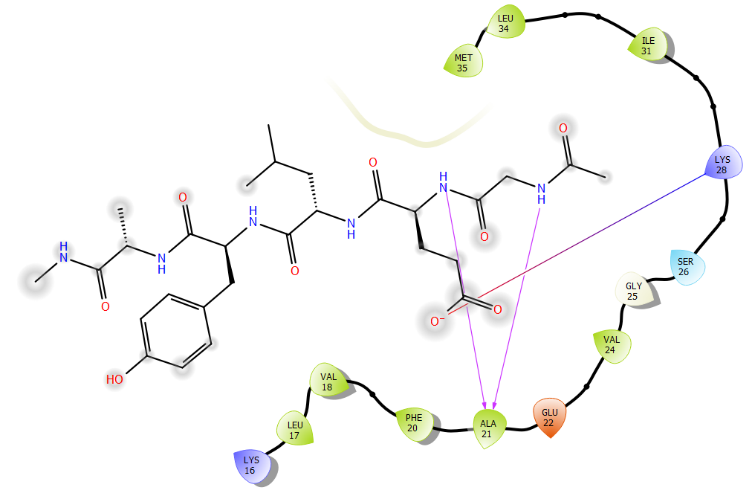

K) **P10**
L) **P9**

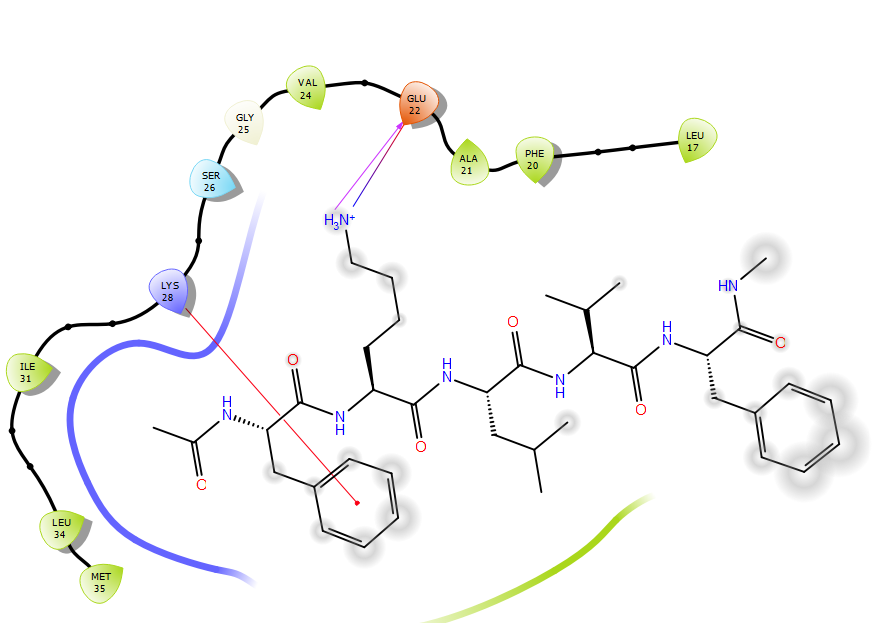

M) **P13**
3. **2D Interaction images of 2MXU with designed peptide inhibitors P1**-**P13 (a**🡪**m)**

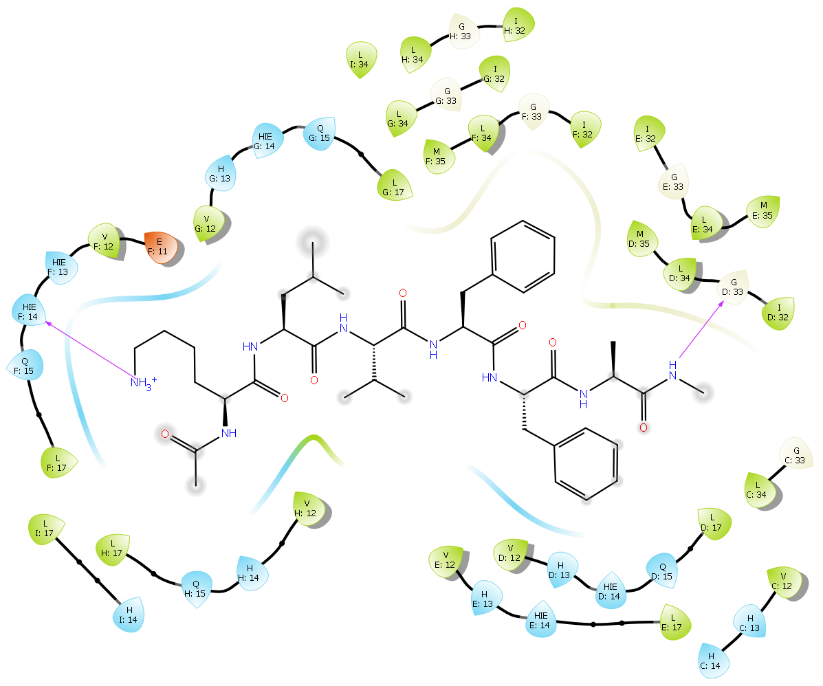

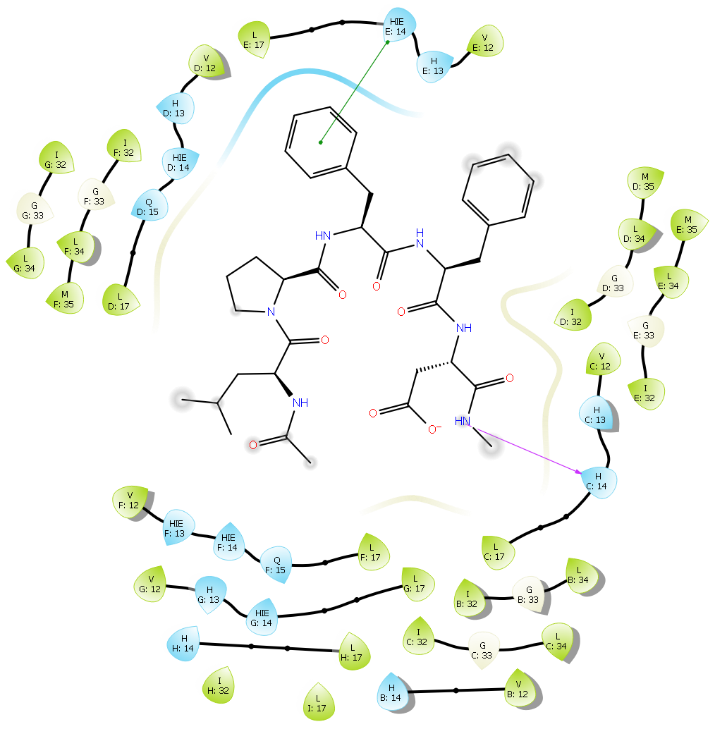

a) **C2**
b) **C1**

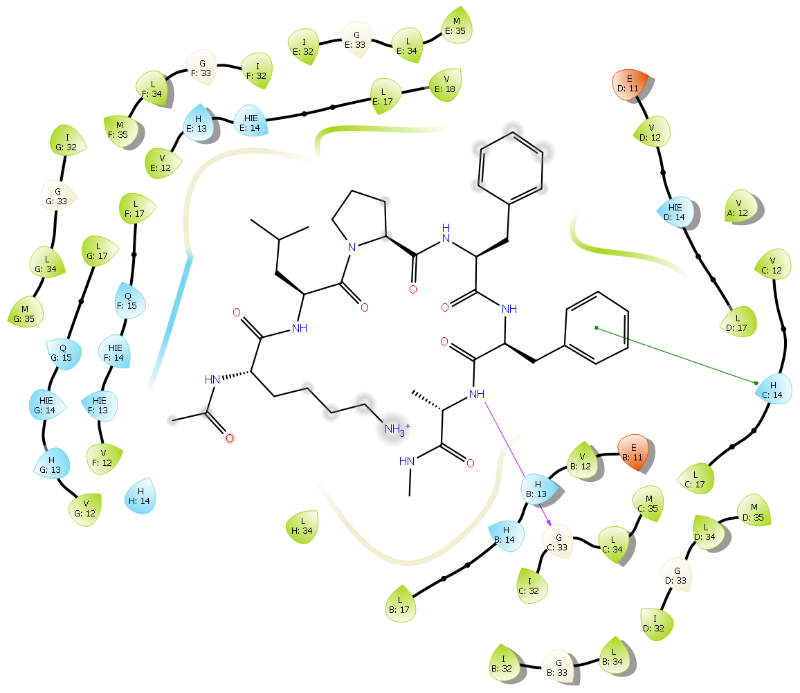

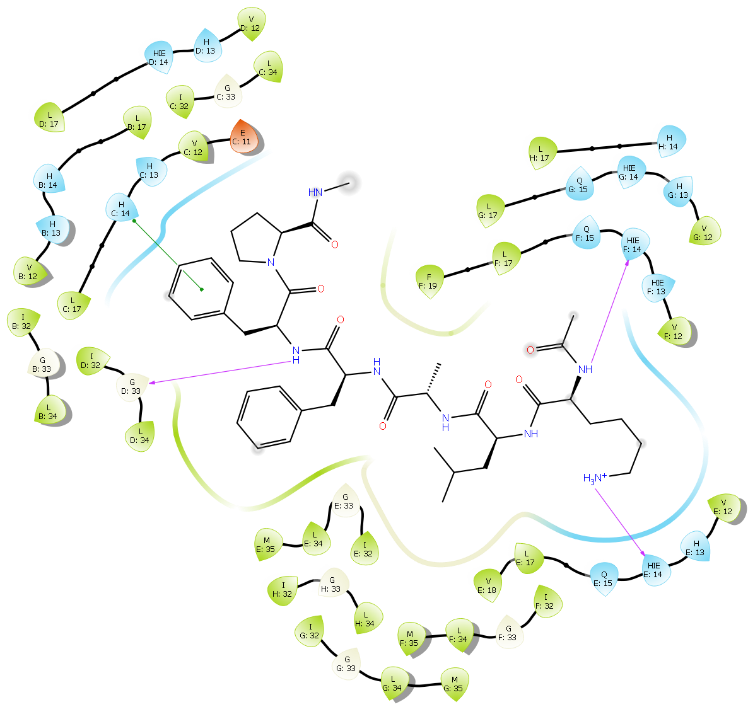

c) **P2**
d) **P1**

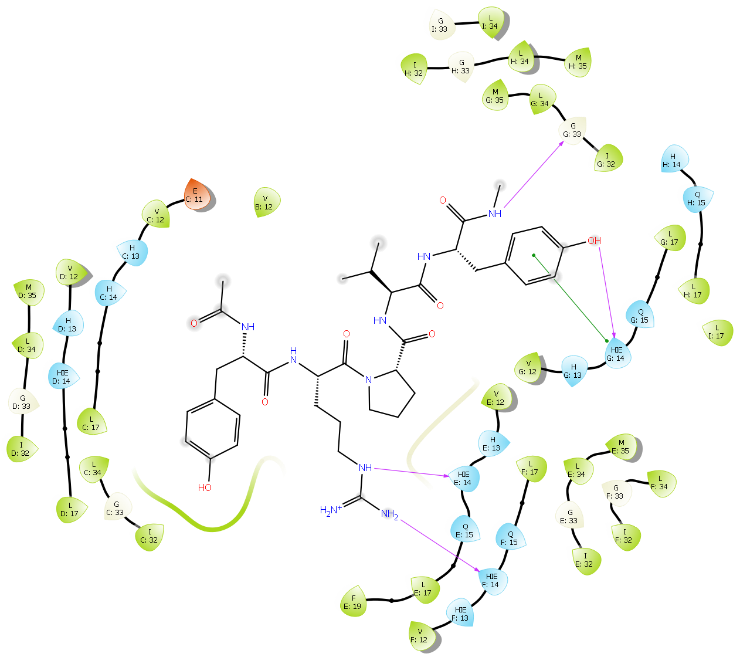

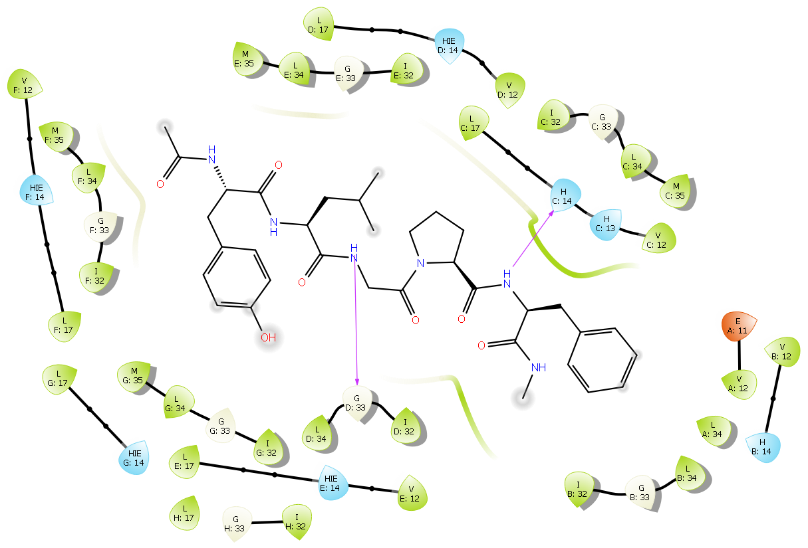

e) **P4**
f) **P3**

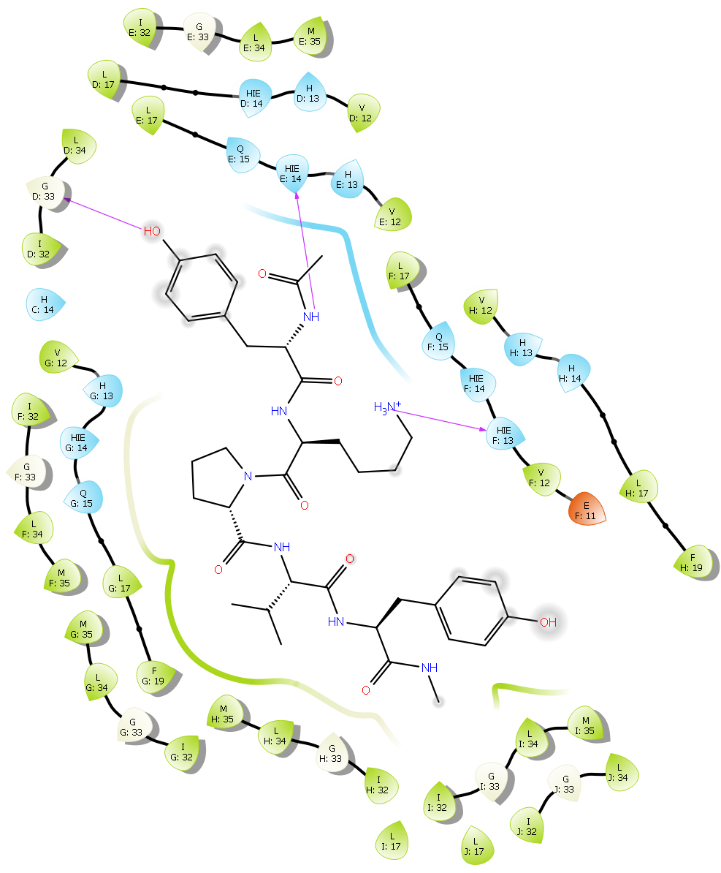

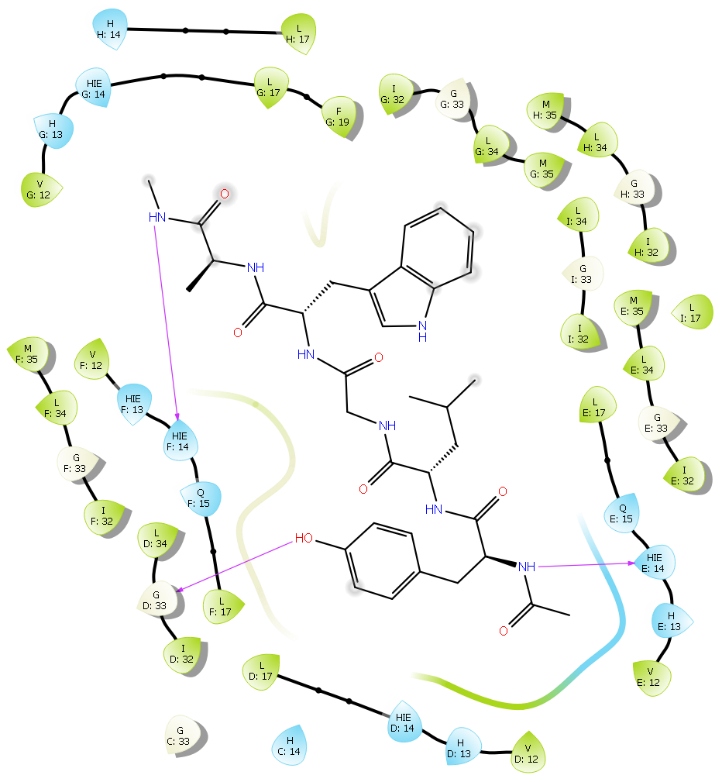

g) **P7**
h) **P5**

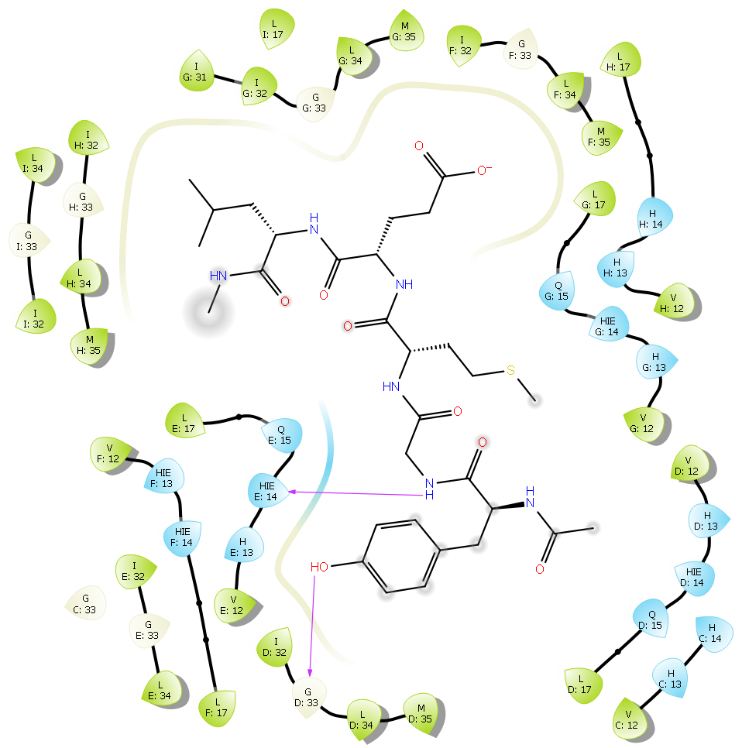

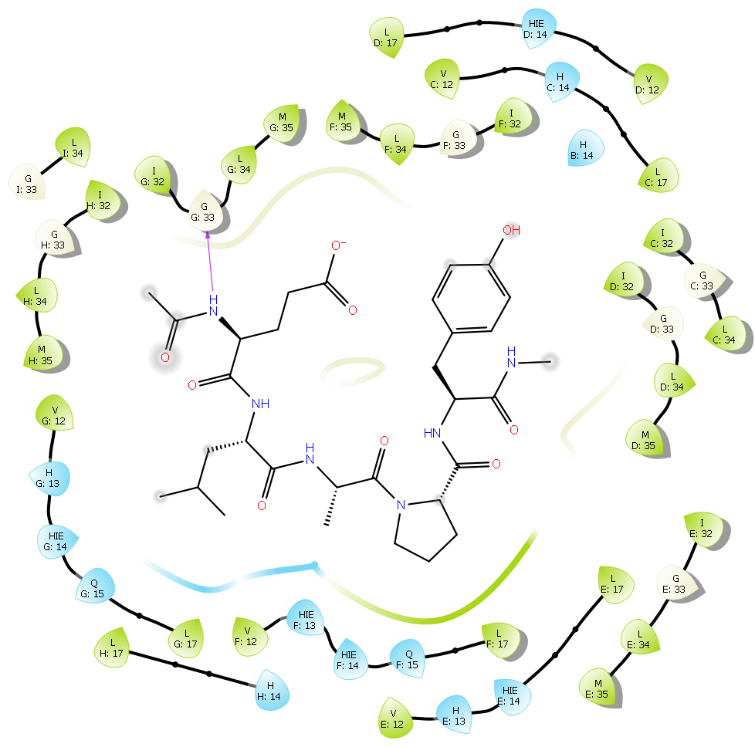

i) **P9**
j) **P8**

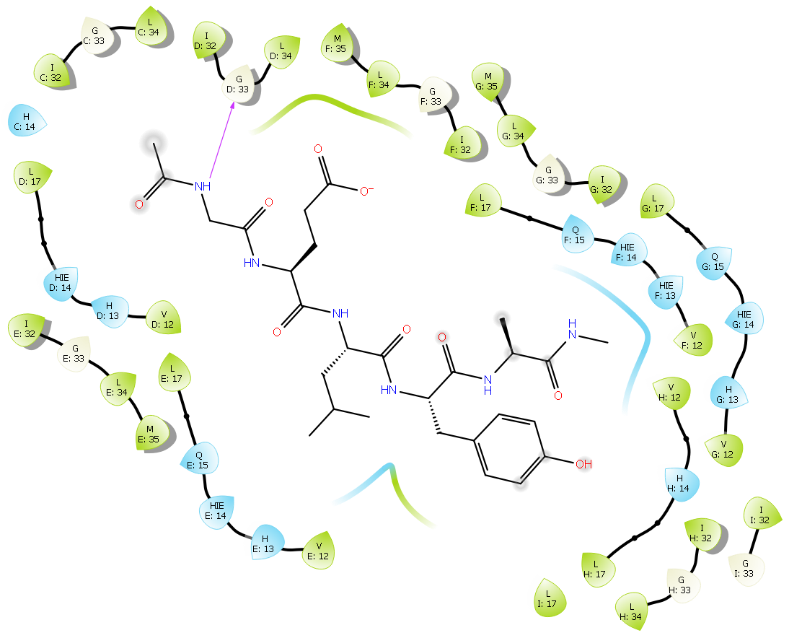

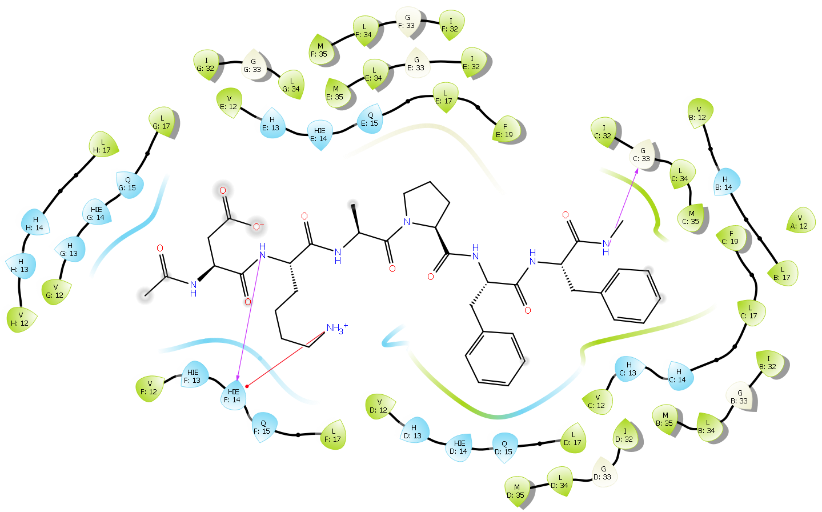

k) **P11**
l) **P10**

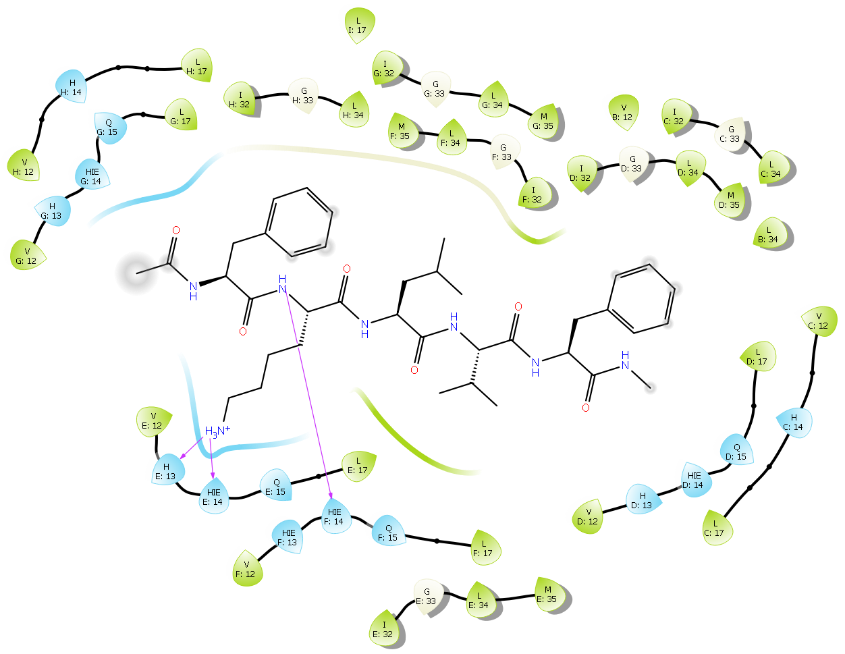

m) **P13**
